# Linker histone H1.0 loads onto nucleosomes through multiple pathways that are facilitated by histone chaperones

**DOI:** 10.1101/2025.02.23.639383

**Authors:** Ehsan Akbari, Nathaniel L. Burge, Michael G. Poirier

## Abstract

Linker histone H1 is an essential chromatin architectural protein that compacts chromatin into transcriptionally silent regions by interacting with nucleosomal and linker DNA, while rapidly exchanging *in vivo*. How H1 targets nucleosomes while being dynamic remains unanswered. Using a single-molecule strategy, we investigated human H1.0 interactions with DNA and nucleosomes. H1.0 directly binds nucleosomes and naked DNA with a preference toward nucleosomes. DNA-bound H1.0 exhibited a range of bound lifetimes with both mobile and immobile states, where the mobile H1.0 did not load onto nucleosomes. The histone chaperone Nap1 facilitated H1.0-nucleosome loading by enabling H1.0 loading through DNA sliding, reducing DNA resident times without impacting nucleosome resident times, increasing mobility along DNA, and targeting H1.0 loading onto the nucleosome dyad. These findings reveal linker histones load onto nucleosomes through multiple distinct mechanisms that are facilitated by chaperones to regulate chromatin accessibility.

## INTRODUCTION

Eukaryotic genomic DNA is organized into chromatin fibers by being repeatedly wrapped into individual nucleosomes, 147bp of DNA wrapped around a histone protein octamer that contains two copies of each core histone: H2A, H2B, H3 and H4 ^1^. The linker histone H1 dynamically regulates chromatin fiber compaction ^2,3^ to control genomic accessibility to transcription and DNA repair machinery ^4,5^. H1 is an abundant chromatin architectural protein that occurs about one H1 per nucleosome in metazoan cells ^6,7^, while H1 expression level is critical for cell viability ^5^. H1 binds within the nucleosome dyad region, stabilizing the DNA wrapped into nucleosomes ^7^ and maintaining chromatin condensation ^8,9^. Interestingly, H1 dynamically exchanges between chromatin fibers ^10–12^, while functioning with other chromatin-modifying complexes to maintain heterochromatic vs. euchromatic states ^4,5^. The molecular mechanism behind linker histone H1 regulation of chromatin states is not fully understood.

All human H1 isoforms ^13,14^ contain a tripartite structure: a globular winged helix domain (WHD) flanked by a short (20-35 residue) disordered N-terminal domain (NTD) and a long (∼100 residue) disordered C-terminal domain (CTD) ^7,15^. Both the WHD and CTD interact with the nucleosome by binding specific nucleosomal DNA regions ^16–18^. The H1 WHD interacts with the DNA open minor groove at the nucleosome dyad and the first 10bp of the linker DNA ^16–18^, while the H1 CTD interacts within ∼20bp of linker DNA, becoming partially ordered upon DNA binding ^19,20^. The positively charged amino acids in H1 CTD neutralize the negatively charged DNA backbone ^21,22^.However, while becoming partially ordered, recent single-molecule evidence ^23,24^ along with NMR analysis^25^ and computational modeling ^26,27^ have revealed that the structure of DNA-bound H1 CTD remains highly dynamic. Collectively, these and other studies have established that H1-WHD and H1-CTD binding to DNA are the primary interactions involved in H1 deposition into chromatin.

Despite interacting with chromatin through high-affinity DNA interactions ^28,29^, H1 localizes onto the nucleosome dyad to form chromatosomes, instead of other nucleosomal DNA or linker DNA regions ^7^. While structural analysis has provided atomic resolution of the H1 WHD within the chromatosome complex ^16,18,30^, how H1 is efficiently targeted to nucleosome dyad while remaining highly dynamic remains an open question. An important regulator of H1 deposition onto chromatin both *in vivo* and *in vitro* is histone chaperones ^31–33^ including Nap1 ^34^. Nap1 has been shown to suppress non-nucleosomal DNA interactions with core histones *in vitro* ^17^, and is required for H1-induced chromatin condensation in cell extracts ^34,35^. Understanding the role of histone chaperones in regulating H1-chromatin interactions is critical for determining H1 regulation of chromatin accessibility.

Here, we report combined single-molecule optical tweezers and fluorescence measurements of H1.0 interaction dynamics to DNA and nucleosomes. We find that most DNA-bound H1.0 is highly mobile, exploring micron distances in seconds by 1D diffusion, while ∼16% is immobile. Both the mobile and immobile fractions of DNA-bound H1.0 remain bound for up to a couple of minutes, where both fractions have fast (sub-second to seconds) and slow (tens of seconds) characteristic resident times. Surprisingly, while we observe direct loading of H1.0 onto nucleosomes, H1.0 alone does not load onto nucleosomes through DNA sliding. However, the histone chaperone Nap1 facilitates efficient H1.0 loading onto nucleosomes by shortening H1.0 resident time on DNA without impacting nucleosome resident time, enhancing H1.0 mobility along DNA, enabling H1.0-nucleosome loading through DNA sliding, and increasing H1.0 loading onto the nucleosome dyad. These measurements reveal that H1.0 interactions with DNA and nucleosomes are highly dynamic and that histone chaperons regulate H1 loading onto nucleosomes through multiple synergistic mechanisms.

## RESULTS

### DNA-bound H1.0 exhibits long residence times with a complex combination of mobile and immobile states

Linker histone H1 simultaneously interacts with both nucleosomal and linker DNA to preferentially bind within the nucleosome dyad region^16^. H1 binding within this region reduces DNA accessibility through increasing chromatin compaction ^36^ and suppressing nucleosome unwrapping ^37,38^. Not surprisingly, because H1 targets chromatin solely through DNA interactions, H1 efficiently binds naked DNA ^28,29^, which is likely important for how H1 is loaded onto chromatin.

To investigate the dynamics of H1 interactions with naked DNA, we relied on a single molecule approach where a 48.5kb lambda DNA molecule is suspended between two streptavidin-coated polystyrene beads by a dual optical trap equipped with confocal fluorescence imaging ^39,40^. This strategy allows for single molecule fluorescence (SMF) detection of fluorophore-labeled H1.0 binding to, translocating along, and dissociating from a single DNA molecule. We focused on H1.0 because it is an extensively studied isoform ^13^ and highly relevant for human disease ^5^. A microfluidic system was utilized to enable stepwise, bottom-up tethering of a single biotin labeled lambda DNA between two streptavidin-coated beads separately held by the two optical traps (**Fig. 1A**, channels 1-3) as described previously ^41^. The formation of a single DNA tether between two beads was confirmed using a worm-like chain (WLC) model of DNA force-extension ^42^. The single tether was then moved into the channel with the imaging buffer containing 3nM H1.0 (**Fig. 1A**, channels 4).

**Figure 1.**
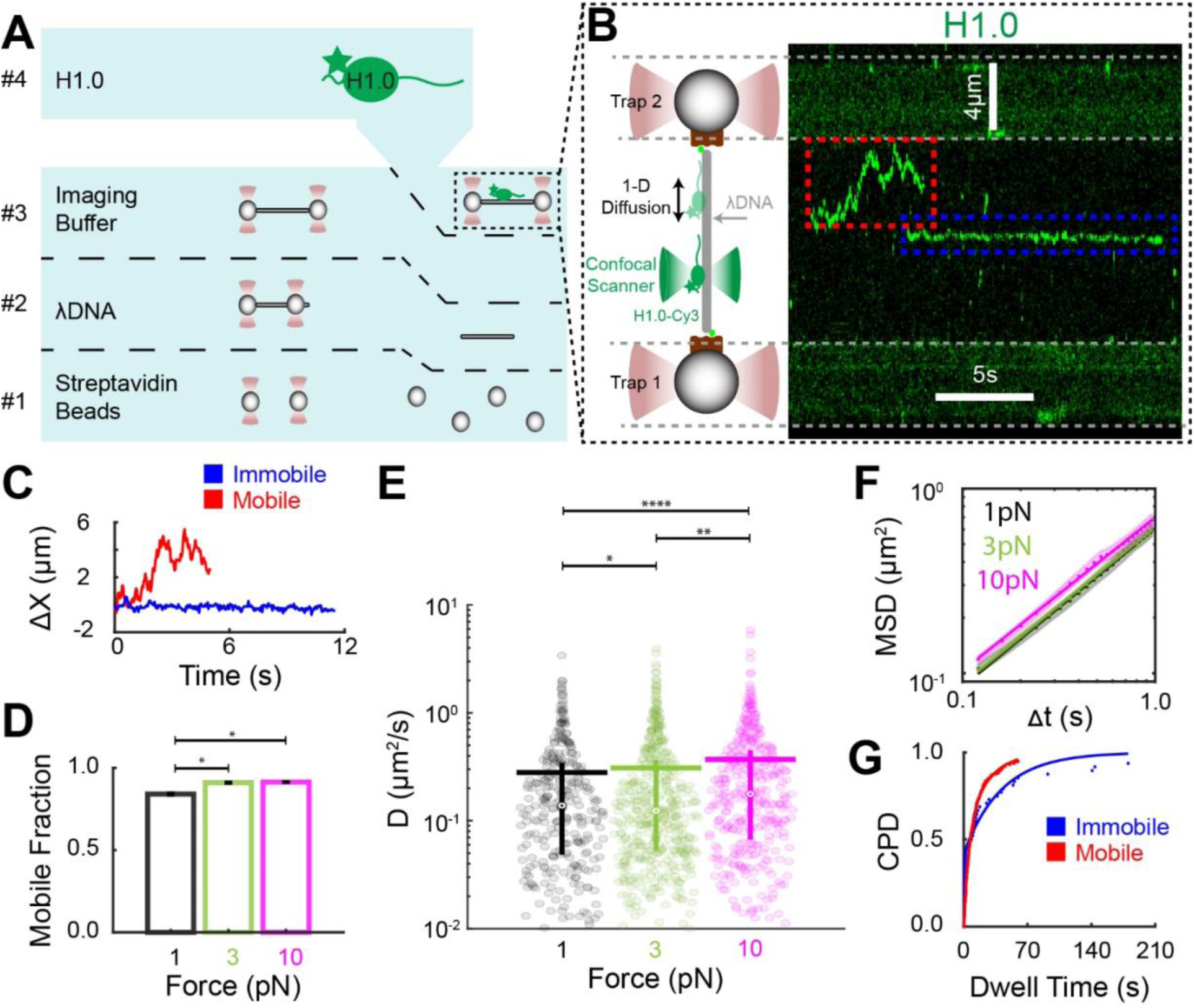
DNA-bound H1.0 is highly dynamic. (**A**) Schematic representation of the microfluidic apparatus utilized for the bottom-up assembly of single molecule H1-dsDNA interaction. (**B**) Schematic of the single-molecule assembly along with a representative kymograph of confocal fluorescent visualization of Cy3-labeled H1.0 during its one-dimensional diffusion along a λDNA tether. The red box indicates a H1.0_mobile_, while the blue dashed box indicates an H1.0_immobile_. (**C**) Net displacement of the H1.0_mobile_ (red) versus H1.0_immobile_ (blue) from the kymograph in (B). (**D**) Fraction of H1.0_mobile_ traces along the λDNA tether held at a constant force of either 1pN (black), 3pN (green), or 10pN (magenta). (**E**) Violin plots of the diffusion constant for each H1.0_mobile_. (**F**) The average mean-squared displacement (MSD) versus time for H1.0_mobile_ traces diffusing along λDNA held at a constant force of either 1pN (black), 3pN (green), or 10pN (magenta). Mechanical force modestly enhances H1.0 diffusion along double-stranded DNA. (**G**) Cumulative probability distribution (CPD) of dwell times observed for mobile (red) and immobile (blue) H1.0 traces under 1pN force. Both dwell time CPDs were best fit to a double-exponential model. * denotes *p*<0.05. ** denotes *p*<0.01. **** denotes *p*<0.0001.

To visualize dynamic binding interactions between DNA and H1.0, we used site-specific Cy3-labeled H1.0 as previously described ^37,43^. The location of H1.0 was determined with SMF confocal line scanning visualized as kymographs of H1.0 interactions along the DNA tether (**Fig. 1B**). The kymographs were recorded while the lambda DNA tether was held under a constant force (1pN, 3pN, or 10pN). These kymographs revealed that Cy3-H1.0 binds to and then remains on the DNA molecule for many seconds before dissociating (**Fig. 1B**). Single-molecule H1.0 traces were quantified to extract the resident time and position as a function of time for each DNA-bound H1.0 molecule. DNA-bound H1.0 exhibited two distinct types of dynamics: mobile (red dash line) and immobile (blue dash line), which were observed for all three constant force values of 1pN (**Fig. 1B-C**), 3pN (**Extended Fig. 1A**) and 10pN (**Extended Fig. 1B**).

To quantitatively determine whether an H1.0 is mobile (H1.0_mobile_) or immobile (H1.0_immobile_), we prepared DNA tethers that contained a sparse number of nucleosomes with Cy5-labeled histone octamer as a reference for immobile DNA-bound protein (**Fig. S1**). Nucleosomes are known to be essentially immobile on the scale of minutes ^44^. Based on the criteria from nucleosome references (see Material and Methods for details), we determined that 84±1, 91±1, and 91.4±0.5 percent of the 308 (1pN), 454 (3pN), and 358 (10pN) tracked H1.0 were diffusing along the DNA tether, respectively (**Fig. 1D**). To quantify the mobility of H1.0_mobile_, we determined the mean squared displacement (MSD) versus time, which fits MSD = 2D t^α^. D is the diffusion constant and α is the exponent of the power law, which is typically close to 1. We find that α is ∼0.8 and independent of applied force (**Fig. 1F**, **Table S1**). However, the distribution of diffusion constants, D, spanned over two orders of magnitude, while the increase in applied force from 1pN to 10 pN resulted in a modest but statistically significant increase in the average D from 0.281±0.002 𝜇m^2^/sec to 0.368±0.003 𝜇m^2^/sec (**Fig. 1E**).

We then investigated the time H1.0 remained bound to DNA with cumulative probability distributions (CPD) of H1.0 dwell times on the DNA tether (**Fig. 1G**). These CPDs were best fit to a double-exponential model for both H1.0_mobile_ and H1.0_immobile_, and for all three forces: 1pN (**Fig. 1G**), 3pN (**Extended Fig. 1C**) and 10pN (**Extended Fig. 1D**). These results imply that DNA-bound H1.0 exhibits two characteristic dwell times largely independent of H1.0 mobility state and the amount of force applied to the DNA. The short multi-second characteristic dwell time (𝜏_fast_) for mobile traces was significantly larger than the sub-second 𝜏_fast_ for the immobile traces at all three forces probed (**Extended Fig. 1E**, **Table S2**). The longer dwell times (𝜏_slow_) were on the scale of 10 seconds or larger, where the mobile and immobile populations were not significantly different for 1 and 3 pN, while at 10 pN, the H1.0_mobile_ 𝜏_slow_ was about 3-fold longer than the H1.0_immobile_ (**Extended Fig. 1E**, **Table S2**). Finally, the fraction of H1.0_mobile_ that dissociates with 𝜏_fast_ (A_fast_) is close to 0.5, implying that H1.0_mobile_ spends similar times bound in the 𝜏_fast_ and 𝜏_slow_ states, while H1.0_immobile_ spends more time bound in 𝜏_fast_ than the 𝜏_slow_ state (**Extended Fig. 1E**, **Table S2)**. Collectively, these findings reveal that H1.0 adopts a range of binding modes to DNA that include highly mobile and immobile states with a wide range of times bound to the DNA.

### H1.0 alone preferentially loads directly onto nucleosomes

Now that H1.0 has been determined to bind to and is mobile on DNA, we investigated how H1.0 loads onto nucleosomes. Properly loaded H1 binds nucleosomes near the dyad symmetry axis, where the H1 WHD interacts with both the DNA near the dyad symmetry axis and the linker DNA as it exits the nucleosome ^18,30^. In contrast, the H1 CTD forms extensive interactions with linker DNA ^16^. Our findings that DNA-bound H1.0 is highly mobile and conformationally dynamic suggest that in addition to H1.0 directly loading onto nucleosomes, H1.0 could undergo 1D diffusion along DNA to find and then load onto the nucleosome near the dyad. Since H1.0 binding within this region allows the WHD to make more DNA contacts than to DNA alone, H1.0 would then preferentially localize within the nucleosome dyad.

To investigate if H1.0 preferentially undergoes direct loading onto nucleosome and if H1.0 utilizes 1-D diffusion along the DNA to load onto nucleosomes, the nucleosomes were reconstituted into biotin-end-labeled lambda DNA, where the nucleosomes were randomly positioned along the tether and contained Cy5-labeled histone octamer at H2A(K119C). This allowed for nucleosome positions to be quantified in kymographs with Cy5 fluorescence. The density of nucleosomes along the DNA tether was prepared with less than 1 nucleosome per 5 kilobase of DNA so that confocal fluorescence scanning could resolve (i) individual nucleosome positions, (ii) sliding of DNA-bound H1.0 between nucleosomes, and (iii) nucleosome-H1.0 interaction events (**Fig. 2A-B**).

**Figure 2.**
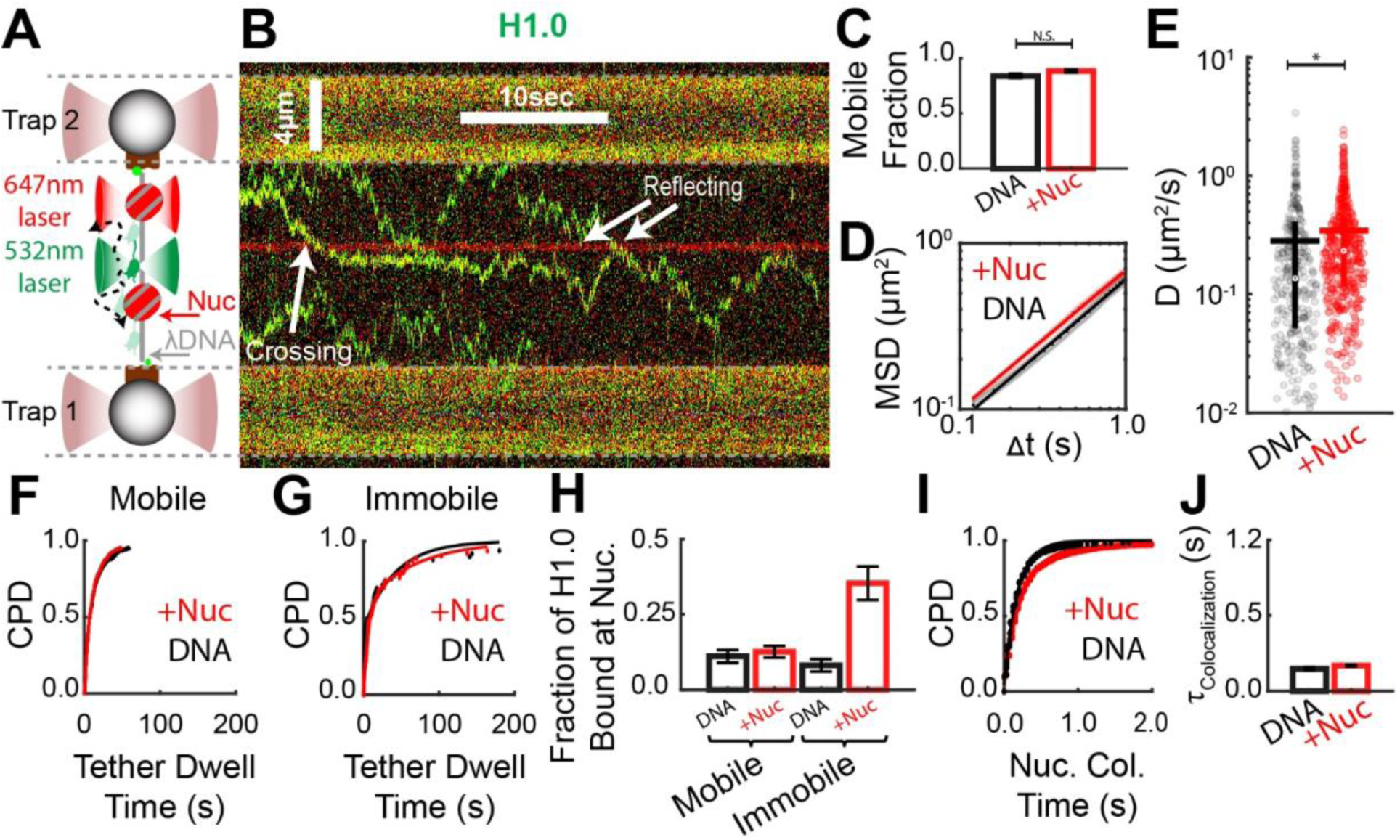
H1.0 directly loads onto nucleosomes but not through DNA sliding. (**A)** Schematic rendering of the single molecule experimental setup to study interactions between nucleosomes and H1.0. (**B)** Representative kymograph of a single nucleosome-containing DNA tether. The red signal indicates the nucleosome (H2A-Cy5) location, and the green signal indicates the H1.0-Cy3 location along the tether. Upon H1.0-nucleosome co-localization, H1.0_mobile_ does not bind to the nucleosome but instead reflected away from the nucleosome. (**C)** Fraction of DNA-bound H1.0 traces that were mobile along DNA-only (black) and nucleosomes-containing (red) tethers. (**D)** Average MSD versus time for H1.0_mobile_ when diffusing along DNA-only (black) and nucleosome-containing (red) tethers. (**E)** Violin plots of diffusion coefficients for H1.0_mobile_ bound to DNA-only (black) and nucleosome-containing (red) tethers. (**F**) and (**G**) Dwell time CPD for H1.0_mobile_ and H1.0_immobile_, respectively, bound to nucleosome-containing (red) and DNA-only (black) tethers. A summary of the fit parameters is in Extended Figure 2A. (**H)** The probability of an H1.0 binding event to a nucleosome-containing tether is colocalized with a nucleosome (+Nuc, red). This is compared to the probability of an H1.0 binding event on a DNA-only tether colocalized at randomly selected points at the same density as the nucleosome density (DNA, black). (**I**) CPD of H1.0_mobile_-nucleosome colocalization events (red) compared to the CPD of H1.0_mobile_ colocalization events at randomly selected points along DNA-only tethers (black). Both CPDs were best fit to a single exponential model. (**J**) A summary of the characteristic dwell times, τ_colocalization_, from the exponential fits of the CPD in (I). The similar τ_colocalization_ suggests that nucleosomes do not reside at nucleosomes longer than a random DNA position. All measurements were done with the tether held at a constant force of 1pN.

Kymographs were acquired with nucleosome-containing lambda DNA that was tethered between two streptavidin-coated beads and subjected to a constant force of 1pN (**Fig. 3A**). 1pN of force was used because it is below the force required to partially unwrap nucleosomes ^45^ and keeps the DNA tether in the image plane for confocal line scanning. Dual fluorescence confocal line scanning was used to simultaneously visualize the positions of Cy5-labeled nucleosomes and DNA-bound Cy3-labeled H1.0 (**Fig. 2A-B**). Upon the addition of 3nM H1.0, we observed tether-bound H1.0, where the fraction of H1.0_mobile_ on nucleosome-containing tethers was not significantly different than on DNA-only tethers (**Fig. 2C, Table S1**). The MSD power law dependence of α for H1.0_mobile_ traces was not impacted by a sparse number of nucleosomes along the tether (**Fig. 2D, Table S1**). However, there was a modest but statistically significant increase in the average diffusion constant of H1.0_mobile_ on nucleosome-containing tethers (**Fig. 2E, Table S1**). This indicates that nucleosomes alone do not significantly impact H1.0 mobility along DNA.

**Figure 3.**
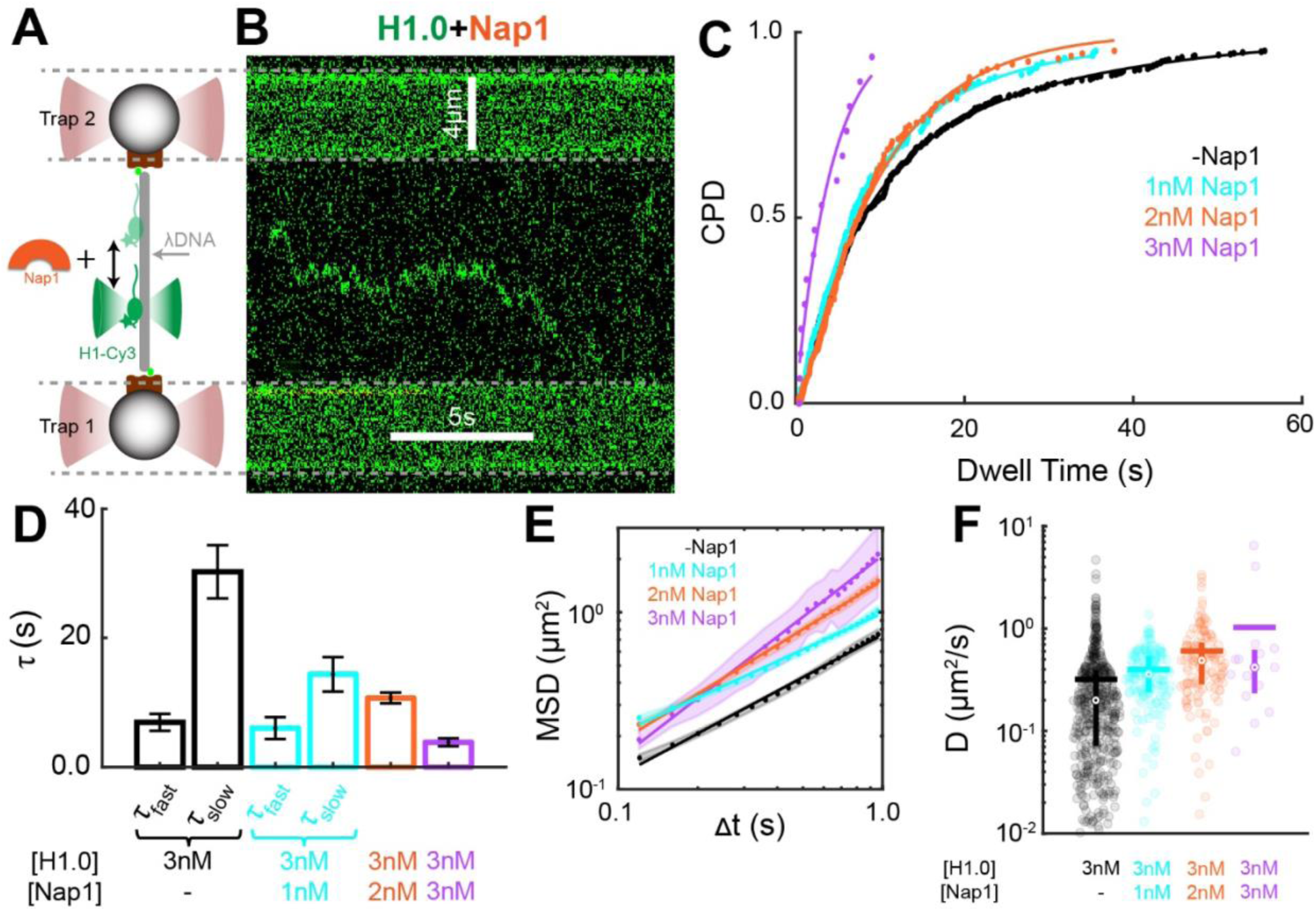
Nap1 reduces H1.0 binding to DNA while enhancing the mobility of DNA-bound H1.0. **(A)** Illustration of the single molecule experimental setup. (**B)** Representative kymograph of a single Cy3-labeled H1.0 (green) diffusing along a DNA-only tether in the presence of Nap1. (**C)** Dwell time CPD of DNA-bound H1.0 in the presence of increasing concentrations of Nap1. The CPDs were best fit with two exponentials for 0 and 1 nM Nap1 and with a single exponential for 2 and 3 nM Nap1. Furthermore, Nap1 systematically reduces the H1.0 dwell time on DNA-only tethers. (**D**) Summary of the characteristic dwell times (τ) from the exponential fits of the CPDs in (C). The double exponential fits provide two characteristic dwell times (τ_fast_ and τ_slow_), while the single exponential fits provide one characteristic dwell time. (**E**) Average MSD versus time for increasing concentrations of Nap1. (**F**) Violin plots of H1.0mobile diffusion coefficients in the presence of increasing concentrations of Nap1. Nap1 systematically increased the mobility of DNA-bound H1.0. All measurements were done with a constant force of 1pN.

We determine the dwell time CPD of mobile and immobile DNA-bound H1.0 on nucleosome-containing tethers. The fit required two exponentials, as was needed for DNA-only tethers (**Fig. 2F-G**). The addition of a sparse number of nucleosomes within the tether did not significantly impact τ_fast_, τ_slow,_ and A_fast_ for the mobile traces (**Extended Fig. 2A, Table S2**). In contrast, for the immobile traces, τ_fast_ and τ_slow_ increased from 0.7±0.2s to 6.6±1.2s and from 48.5±9.2s to 79.0±14.9s due to sparsely positioned nucleosomes, respectively (**Extended Fig. 2A, Table S2**). Meanwhile, the addition of a sparse number of nucleosomes did not significantly change A_fast_ for the immobile traces (**Extended Fig. 2A, Table S2**).

These findings suggest that while the presence of nucleosomes did not impact the resident time of H1.0_mobile_ on the tether, nucleosomes do impact the resident time of H1.0_immobile_. To further investigate this, we separately quantified the fraction of H1.0_mobile_ and H1.0_immobile_ that colocalized with nucleosomes when first binding nucleosome containing tethers (**Fig. S2**) as compared to binding at randomly chosen positions along DNA-only tethers. The number of random positions was set to 4, the average number of nucleosomes per tether. We found that H1.0_mobile_ did not bind directly to nucleosomes more often than to random DNA positions (**Fig. 2H**). However, H1.0_immobile_ bound to nucleosomes about 4-fold more frequently than to random DNA positions (**Fig. 2H**). These results indicate that H1.0_immobile_ preferentially loads directly onto nucleosomes more often than random binding to DNA alone, resulting in the increased fraction and resident time of H1.0_immobile_.

### H1.0 alone does not load onto nucleosomes through DNA sliding and instead reflects away from nucleosomes

Interestingly, visual inspection of the kymographs indicated that upon encountering a nucleosome, H1.0_mobile_ reflected away from the nucleosome, while occasionally crossing over the nucleosome (**Fig. 2B, Fig. S3**). Furthermore, we did not often observe H1.0_mobile_ that encountered a nucleosome through 1D diffusion remaining bound to the nucleosome (**Fig. S3**). To quantitatively analyze the interactions between mobile DNA-bound H1.0 and nucleosomes, we defined a colocalization event when the position of H1.0 was within ±200nm of the nucleosome position. We then determined the CPD of the nucleosome colocalization dwell times and found there was essentially no difference between H1.0 colocalized at either a nucleosome or a random location along a DNA-only tether (**Fig. 2I, J**). These results indicate that mobile DNA-bound H1.0 that encounter nucleosomes through DNA sliding do not preferentially reside at nucleosome locations.

During each colocalization event, four outcomes were possible: binding, reflecting, crossing, and dissociating (**Extended Fig. 2B-C**). The CPD of colocalization times between H1.0 and nucleosomes revealed a ∼0.2s characteristic colocalization time (τ_colocalization_, **Fig. 2J**). Hence, a colocalization event was considered a binding event if the H1.0 and nucleosome were colocalized for more than 0.4s (twice the characteristic dwell time for nucleosome colocalization). If a colocalization event is not a binding event, it was considered as a crossing, reflecting, or dissociating event based on whether the sliding H1.0 maintained direction, reversed direction, or dissociated, respectively. To control for the mobility of H1.0 on DNA-only tethers, colocalization between H1.0 and randomly selected points along the lambda tether without nucleosomes was determined (**Fig. 1**, 1pN of force). The binding probability of H1.0 to nucleosomes was not statistically different than to random points along a DNA-only tether (**Extended Fig. 2D, Table S3**). Instead, upon colocalization between H1.0 and nucleosomes, the H1.0 molecule exhibited a higher likelihood of reflecting off a nucleosome. (**Extended Fig. 2D, Table S3**). These results suggest that H1.0 alone is not able to efficiently load onto a nucleosome if bound first to linker DNA. Furthermore, nucleosomes constrain mobile DNA-bound H1.0, limiting the distance it can travel by 1D diffusion.

### Nap1 efficiently regulates the lifetime and mobility of DNA-bound H1.0

The histone chaperone Nap1 is reported to regulate linker histone H1 interactions with chromatin in vivo ^34,35,38^, and directly interact with linker histones in vitro ^46,47^. Based on our findings that DNA-bound H1.0 is conformationally dynamic and does not efficiently load onto nucleosomes through DNA sliding, we considered the possibility that Nap1 regulates H1.0 interactions with chromatin in part by influencing the lifetimes and mobility of DNA-bound H1.0.

To investigate the impact of Nap1 on H1.0-DNA dynamics, we acquired kymographs of DNA-bound H1.0 with 1pN of constant force and increasing concentrations of Nap1 (**Fig. 3A-B** and **Supplementary Fig. S4**). We observed that at sub-stoichiometric concentrations of Nap1 (< 3nM), H1.0 continued to bind DNA, while Nap1 concentrations above 3nM largely abolished H1.0 binding to DNA (**Fig. S4**). We therefore focused on Nap1 concentrations below 3nM. Interestingly, at these concentrations of Nap1, we did not observe an immobile fraction of DNA-bound H1.0. Therefore, we used the mobile fraction of DNA-bound H1.0 without Nap1 as the comparison group for the mobility and lifetime measurements with Nap1.

We first focused on the impact of Nap1 on the mobility of DNA-bound H1.0. The average time dependence of the MSD over a 1s time window fit a power law of α ∼ 1 as the Nap1 concentration was increased (**Fig. 3E**, **Table S1**). In addition, the average diffusion constant of H1.0_mobile_ without Nap1 (D=0.281±0.002 𝜇m^2^/sec) increased to 0.40±0.02 𝜇m^2^/s, 0.60±0.06 𝜇m^2^/s, and 0.95±0.29 𝜇m^2^/s, with 1nM, 2nM, and 3nM of Nap1, respectively (**Figure 3F**, **Table S1**). These results reveal that Nap1 significantly enhances unconstrained random walk diffusion of DNA-bound H1.0.

In addition, we quantified the dwell time of each DNA-H1.0 binding event and then fit the CPD of these dwell times to exponential functions (**Fig. 3C**). We find that the range of DNA-bound H1.0 dwell times systematically decreased with increasing Nap1 (**Fig. 3C**). In the presence of 1nM of Nap1, 2 exponential functions were required to fit the H1.0 dwell time CPD where the 𝜏_fast_ did not change, while 𝜏_slow_ decreased from 23.2±1.7s to 14.4±2.7 s **(Fig. 3C-D, Table S2**). Furthermore, the fraction of fast transitions, A_fast_, increased from A_fast_ = 55±9 % to A_fast_ = 70±20 %. In contrast, the dwell time CPD of DNA-bound H1.0 with 2nM and 3nM of Nap1 only required a single exponential with a characteristic dwell time of 10.7±0.8 s and 3.8±0.6 s, respectively **(Fig. 4C-D, Table S2**). These results indicate that sub-stoichiometric concentrations of Nap1 not only prevent H1.0 from residing on DNA in the immobile state but also suppress the long-lived bound states of H1.0_mobile_. Collectively, these studies reveal that Nap1 influences the occupancy and mobility of DNA-bound H1.0, suggesting Nap1 can regulate H1.0 binding to chromatin via controlling H1.0 interactions with linker DNA.

**Figure 4.**
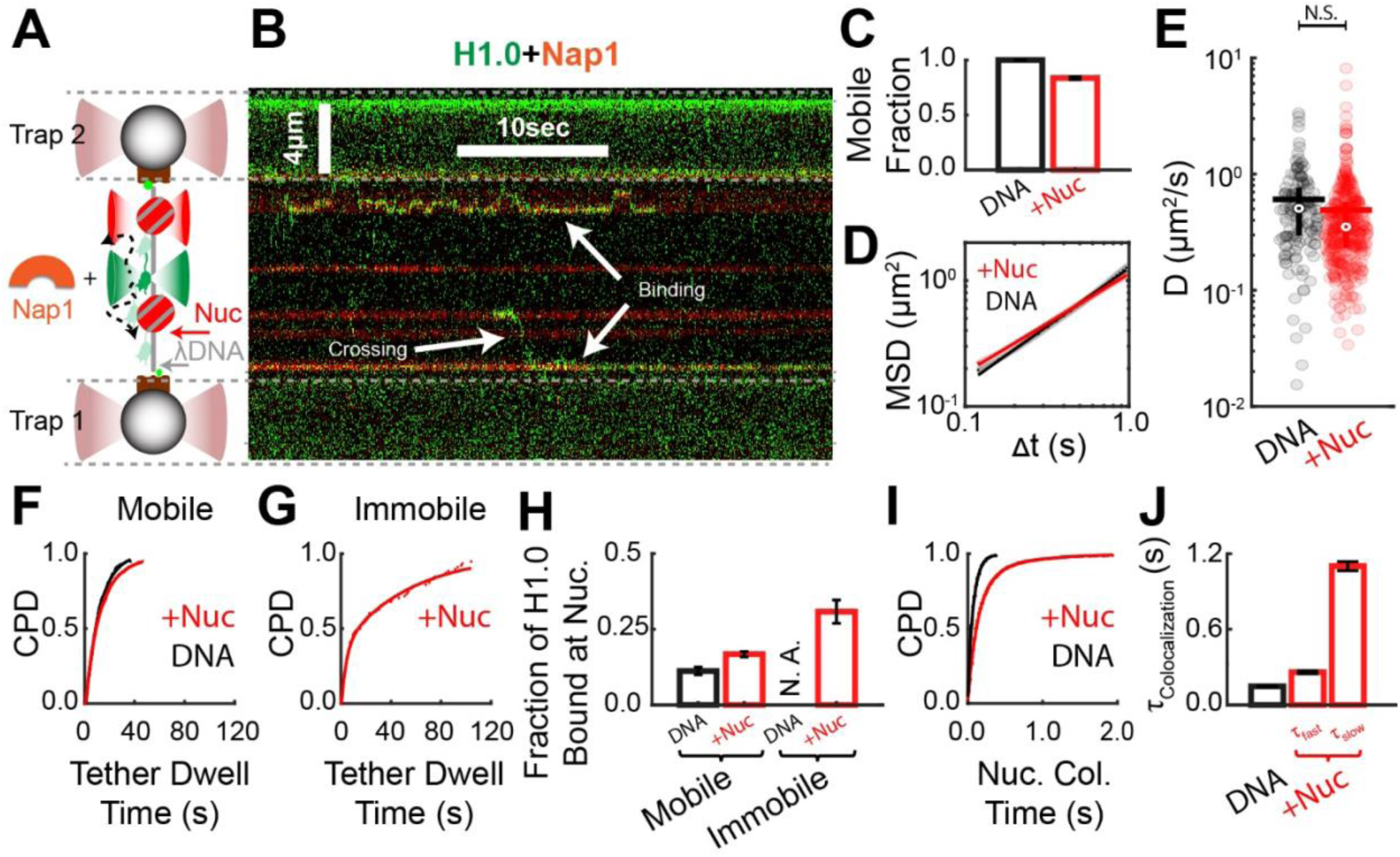
Nap1 facilitates the efficient loading of H1.0 onto nucleosomes. **A)** Illustration of the single molecule experiment. (**B**) Representative kymograph of H1.0 (green) bound to a nucleosome-containing tether in the presence of Nap1. Red indicates the locations of the nucleosomes along the DNA tether. (**C**) Fraction of mobile DNA-bound H1.0 in the presence of Nap1 when bound to DNA-only (black) and nucleosome-containing (red) tethers. (**D**) Average MSD versus time of H1.0_mobile_ traces when diffusing along nucleosome-containing (red) and DNA-only (black) tethers. (**E**) Violin plots of diffusion coefficients for H1.0_mobile_ along DNA-only (black) and nucleosome-containing (red) tethers in the presence of Nap1. (**F**) and (**G**) Dwell time CPD for H1.0_mobile_ and H1.0_immobile_, respectively, bound to DNA-only (black) and nucleosome-containing (red) tethers in the presence of Nap1. Nap1 eliminated the presence of H1.0_immobile_ on DNA-only tethers, so it is not included in (G). A summary of the fit parameters is in Extended Figure 2A. (**H**) In the presence of Nap1, the probability of an H1.0 binding event to a nucleosome-containing tether is colocalized with a nucleosome (+Nuc, red). This is compared to the probability of an H1.0 binding event to a DNA-only tether colocalized with randomly selected points at the same density as the nucleosome density (DNA, black). (**I**) In the presence of Nap1, the CPD of H1.0mobile-nucleosome colocalization events (red) compared to the CPD of H1.0_mobile_ colocalization events at randomly selected points along DNA-only tethers (black). The CPD at random positions on DNA-only tethers was best fit to a single exponential model, while the CPD at nucleosomes was best fit to a double exponential. (**J**) A summary of the characteristic dwell times, τ_colocalization_, from the exponential fits of the CPDs in (I). The addition of Nap1 resulted in a second characteristic time, τ_slow_, suggesting that Nap1 facilitates stable loading of H1.0 onto nucleosomes. All measurements were done with the tether held at a constant force of 1pN and with 2nM of Nap1.

### Nap1 enhances H1.0 loading efficiency onto nucleosomes by eliminating immobile DNA-bound H1.0 while not impacting direct loading of H1.0 onto nucleosomes

Given our observation that Nap1 is a strong modulator of H1-DNA binding interactions, combined with reports that Nap1 regulates H1.0-nucleosome interactions *in vitro* and *in vivo* ^35,47^, we investigated the possibility that Nap1 influences H1.0 loading efficiency onto nucleosomes. To do this, 2nM of Nap1 was included in kymograph measurements of Cy3-labeled H1.0 interacting with DNA tethers containing Cy5-labeled nucleosomes while being held at a constant force of 1pN (**Fig. 4A, B**). The kymograph measurements were then quantified to determine the impact of Nap1 on the fraction of H1.0_mobile_ and the CPD of H1.0 bound dwell times. We first determined the fraction of H1.0_mobile_ on nucleosome-containing DNA in the presence of Nap1. We observed that 84±1% of all bound H1.0 was mobile (**Fig. 4C**). This is in stark contrast to the impact of 2nM of Nap1 on H1.0 bound to DNA-only tethers, where all H1.0_immobile_ was eliminated, resulting in 100% H1.0_mobile_.

We then investigated the impact of Nap1 on the time H1.0 remains bound to nucleosome-containing DNA tethers by determining the dwell time CPD of H1.0 bound in the presence of Nap1, and compared this to both the dwell time CPD of H1.0 bound to DNA-only tethers with Nap1 (**Fig. 4F-G**) and the dwell time CPD of H1.0 on nucleosome-containing tethers without Nap1 (**Fig. 2F-G**). The CPD for H1.0_mobile_ bound to nucleosome-containing tethers in the presence of Nap1 fit best to a double exponential with a 𝜏_fast_ = 9.1±2.0s and a 𝜏_slow_ = 20.0±4.1s (**Extended Fig. 4A, Table S2**). This 𝜏_fast_ is similar to the 𝜏 = 10.7±0.8 s for H1.0_mobile_ on DNA-only tethers in the presence of Nap1 (**Table S2**. However, H1.0_mobile_ bound to DNA-only tethers did not have a second slower 𝜏 (**Fig. 3D**). In addition, a comparison between the dwell time CPDs of H1.0_mobile_ on nucleosome-containing tethers with Nap1 (**Fig. 4F**) and without Nap1 (**Fig. 2F**) revealed that both required two exponentials with similar 𝜏_fast_ and 𝜏_slow_. Combined, these results indicate the dwell time distribution of H1.0_mobile_ on nucleosome-containing tethers is not drastically impacted by the sub-stoichiometric Nap1 concentration of 2nM.

We also determined the dwell time CPD for H1.0_immobile_ on nucleosome-containing tethers in the presence of Nap1 and found it fit best with 2 exponentials (𝜏_fast_ , 𝜏_fast_ and A_fast_ of 4.2±1.4s, 55.5±4.0s, and 48±7%, **Fig. 4G and Extended Fig. 4A**). This is in stark contrast to our observation that Nap1 eliminates H1.0_immobile_ on DNA-only tethers (**Fig. 4C**). Interestingly, these dwell times were similar to H1.0_immobile_ on nucleosome-containing tethers in the absence of Nap1 (**Extended Fig. 2A**). These results reveal that while the presence of nucleosomes does not dramatically impact the influence of Nap1 on H1.0_mobile_, nucleosomes were required for a fraction of H1.0_immobile_ to form in the presence of Nap1 concentrations that eliminate H1.0_immobile_ on DNA-only tethers.

Based on our observation that H1.0_immobile_ preferentially binds nucleosomes directly without first DNA sliding (**Fig. 2H**), we investigated the impact of Nap1 on the direct binding of H1.0 to nucleosomes. In the presence of 2nM Nap1, we compared the direct binding of H1.0 onto nucleosomes (**Fig. S5**) relative to binding random positions within a DNA-only tether where the number of random positions is the same as the number of nucleosomes per tether. As we observed without Nap1, we found that the fraction of H1.0_mobile_ colocalized with nucleosomes when first loaded onto the tether was the same as binding to random DNA positions (**Fig. 4H**). However, ∼30% of H1.0_immobile_ directly loaded onto nucleosomes, which is similar to the fraction of H1.0_immobile_ that directly loaded onto nucleosomes without Nap1. This is in stark contrast to our observation that Nap1 fully suppressed H1.0_immobile_ on DNA-only tethers (**Fig. 4H**). These results imply that Nap1 enhances H1.0 targeting to nucleosomes in part by suppressing H1.0-DNA binding without impacting H1.0 loading onto nucleosomes directly.

### Nap1 enables the loading of mobile DNA-bound H1.0 onto nucleosomes through DNA sliding

Given that Nap1 eliminated H1.0_immobile_ and increased the H1.0 mobility on DNA-only tethers (**Fig. 3**), we investigated the mobility of H1.0 on nucleosome-containing tethers. To do this, we analyzed kymographs of Cy3-labeled H1.0 interacting with nucleosome-containing tethers that are held at a constant force of 1pN and in the presence of 2nM Nap1. We first determined the average MSD over 1s, which revealed that Nap1 induced a modest but not statistically relevant reduction in the diffusion constant for the H1.0_mobile_ fraction on nucleosome-containing tethers as compared to DNA-only tethers (**Fig. 4E, Table S1**). This is in contrast to the impact of Nap1 on DNA-only tethers, which results in a dramatic increase in the H1.0 diffusion constant (**Fig. 3F**). This suggests that in the presence of Nap1, H1.0 interactions with nucleosomes slow H1.0 diffusion. Interestingly, the presence of nucleosomes reduced the power law dependence of the MSD on time from α = 0.92±0.01 to α = 0.79±0.01 (**Fig. 4D**, **Table S1**), which indicates the presence of nucleosomes constrains the distance H1.0 explores by diffusion.

Visual inspection of the kymographs of Cy3-labeled H1.0 bound to DNA tethers containing Cy5-labeled nucleosomes revealed that, in the presence of Nap1, H1.0 co-localizes to nucleosomes for many seconds and then transitions onto linker DNA where it is mobile until it co-localizes at another nucleosome, or until it dissociates from the tether (**Fig. 4B, Fig. S6**). To quantify this, we determined the CPD for H1.0-nucleosome co-localization times when 2nM Nap1 is included (**Fig. 4I**). In the presence of Nap1, the dwell-time CPD for H1.0 co-localization at nucleosomes was significantly extended compared to co-localization at randomly selected positions along DNA-only tethers (**Fig. 4I**). The nucleosome co-localization dwell-time CPD fit best to a double exponential with a 𝜏_fast_ = 0.30±0.01 s and a 𝜏_slow_ = 1.10±0.04 s while the dwell-time CPD at randomly selected DNA positions fit to a single exponential with a 𝜏 = 0.20±0.01 s (**Fig. 4J**).

We then determined for each colocalization event if H1.0 bound to, crossed over, reflected off, or dissociated from the nucleosome (**Extended Fig. 4C**). Using the criteria that H1.0-nucleosome co-localization events longer than 0.4s were defined as binding events, we determined that the presence of Nap1 increased H1.0-nucleosome binding probability by ∼3-fold as compared to randomly selected positions along DNA-only tethers (**Extended Figures 4B-D, Table S3**). Nap1 did not significantly impact the observation that H1.0 was more likely to reflect away from nucleosomes than cross over nucleosomes among the fraction of H1.0-nucleosome colocalization events that did not result in binding (compare **Extended Fig. 4D** to **Extended Fig. 2D, Table S3**). However, the DNA-only control indicates that Nap1 causes H1.0 to be biased toward continuing in the same direction instead of being equally likely to continue in the same direction or reverse directions (compare **Extended Fig. 4D** to **Extended Fig. 2D, Table S3**). In combination, these results reveal that Nap1 facilitates the loading of H1.0_mobile_ onto nucleosomes in part by both increasing the mobility of DNA-bound H1.0 and enabling H1.0 loading onto nucleosomes through DNA sliding.

### Nap1 facilitates H1.0 loading into the dyad region of nucleosomes

While these co-localization measurements show Nap1 significantly enhances H1.0 loading onto nucleosomes, they do not determine if H1.0 is bound in the nucleosome dyad region as resolved in structural studies ^16,30^. To investigate if H1.0 is localized near the nucleosome dyad during a binding event, we utilized Förster Resonance Energy Transfer (FRET) efficiency measurements between Cy3-labeled H1.0(S22C) and Cy5-labeled histone octamer at H2A(K119C) (**Fig. 5A-B**). In a previous study, we demonstrated that H1.0 binding to the nucleosome dyad region resulted in a significant increase in FRET efficiency ^37^. The H1.0 kymographs were analyzed to quantify the Cy5 photon count (acceptor signal) and Cy3 photon count (donor signal) during binding events between H1.0 and nucleosomes. To estimate the FRET efficiency, the acceptor signal divided by the total detected signal (donor signal + acceptor signal) was quantified. Nucleosome-bound H1.0 exhibited both low and high FRET states, suggesting that not all binding events involved H1.0 integrating near the nucleosome dyad (**Fig. 5C-D**). We determined the average FRET efficiency of each binding interaction for 267 binding events with Nap1 and 64 binding events without Nap1. Furthermore, we separated binding events into binding through direct nucleosome loading versus binding through DNA sliding (**Fig. 5E**). While there was a broad distribution of FRET efficiencies with and without Nap1, the average FRET efficiency of H1.0 that directly loaded onto nucleosomes with Nap1 was 0.73±0.02, which is significantly higher than 0.37±0.04, the FRET efficiency without Nap1 (**Extended Fig. 5D**). Similarly, the average FRET efficiency of H1.0 that loaded onto nucleosomes through DNA sliding with Nap1 was 0.68±0.01, which is also significantly higher than 0.35±0.01, the FRET efficiency without Nap1 (**Fig. 5E**). Overall, these results reveal that in addition to Nap1 enhancing H1.0 targeting to nucleosomes, Nap1 helps H1.0 load into the dyad region of the nucleosome both when loaded directly and when loaded through DNA sliding.

**Figure 5.**
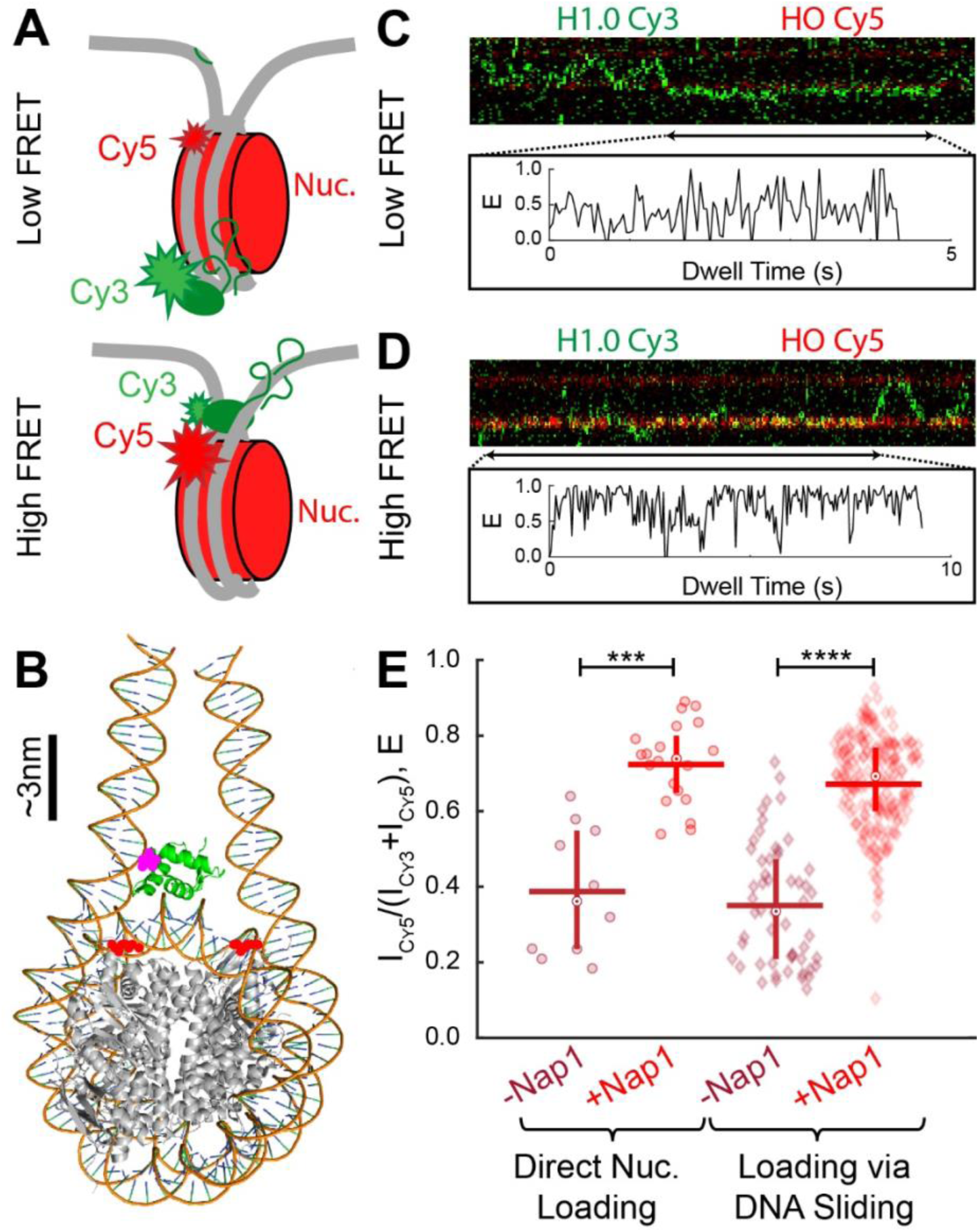
Nap1 facilitates H1.0 loading near the nucleosome dyad. (**A**) Illustration of the FRET construct for detecting dyad binding of H1.0 within nucleosomes. (**B**) Structural model of human H1.0 (green, PDB ID 7K5X) bound to the nucleosome with the locations of Cy3 on H1.0 S22C (magenta) and Cy5 on H2A K119C (red). The histone octamer is depicted in gray, while the DNA backbone is depicted in light brown. (**C**) and (**D**) Representative traces of nucleosome-bound H1.0 exhibiting low FRET and high FRET, respectively. (**E**) Violin plots of the average FRET efficiency during H1.0-Nucleosome colocalization events with (red) and without (dark red) Nap1. Direct nucleosome loading are events where the H1.0-nucleosome colocalization occurred when H1.0 first loaded onto nucleosome-containing tethers. Loading via DNA sliding are events where H1.0-nucleosome colocalization occurred by H1.0 sliding. All measurements were done with a constant force of 1pN.

## DISCUSSION

Using a single molecule strategy that combines dual optical trapping with confocal fluorescence microscopy, we directly visualized and quantified the dynamics of H1.0 interactions with DNA and H1.0 loading onto nucleosomes. We find that H1.0 alone preferentially targets nucleosomes over naked DNA by about four-fold (**Fig. 6A**), which is likely one pathway for nucleosome loading *in vivo*. However, while H1.0 does bind DNA strongly, where ∼84% rapidly diffuses along DNA and encounters nucleosomes, H1.0 alone is unable to load onto nucleosomes through DNA sliding (**Fig. 6B**). Instead, it preferentially reflects away from the nucleosome. These results suggest H1.0 does not efficiently load onto nucleosomes alone. However, the histone chaperone Nap1 helps H1.0 load onto nucleosomes by four separate mechanisms. (i) Nap1 suppresses H1.0 binding to DNA without impacting H1.0 direct binding to nucleosomes (**Fig. 6C-D**). (ii) Nap1 enhances H1.0 mobility along DNA, allowing H1.0 to diffuse kilobases and repeatedly encounter nucleosomes (**Fig. 6D**). (iii) Nap1 allows H1.0 loading onto nucleosomes through DNA sliding, providing an additional pathway for nucleosome loading (**Fig. 6D**). (iv) Nap1 increases the probability that H1.0 binds within the nucleosome dyad so that it is properly loaded (**Fig. 5E**). These mechanisms are critical for H1.0 to suppress DNA accessibility through reducing nucleosome partial unwrapping and enhancing chromatin compaction.

**Figure 6.**
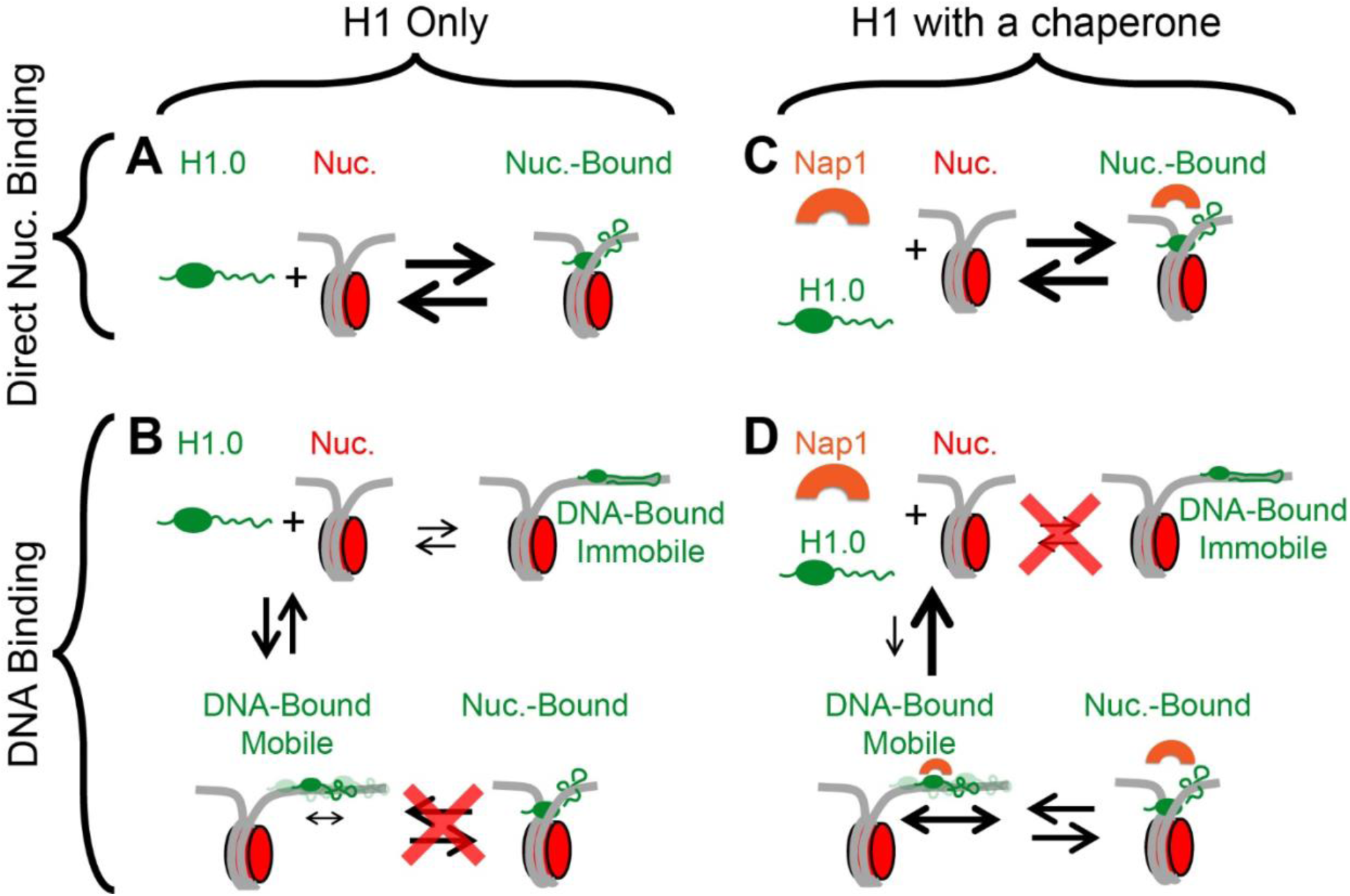
Model of H1.0 loading onto nucleosomes and the impact of Nap1 in facilitating the loading modes of H1.0. (**A)** In the absence of the H1 chaperone Nap1, H1.0 can directly load onto nucleosomes. (**B**) In the absence of Nap1, H1.0 directly binds DNA where the majority fraction is mobile on the DNA but unable to load onto nucleosomes. A minor fraction is immobile and, therefore, unable to load onto nucleosomes. (**C**) H1.0 direct loading onto nucleosomes is unaffected by Nap1. (**D**) H1.0 loading onto DNA in the immobile state is blocked by Nap1. H1.0 still loads onto DNA in a mobile state, but Nap1 lowers the binding rate and increases the dissociation rate. However, Nap1 increases the 1D mobility of H1.0 and facilitates H1.0 loading onto nucleosomes through DNA sliding.

Importantly, these findings reveal that H1.0 in combination with a histone chaperone uses multiple pathways for H1.0 targeting and loading onto nucleosomes. While there has been significant interest in the precise interactions H1 makes to both dyad and linker DNA, which results in on-dyad vs. off-dyad binding ^48^, less is known about the pathway(s) for H1 loading onto nucleosomes. Previous work has proposed that H1 interactions with linker DNA are important for linker histone loading ^12,49^. Our studies reveal that linker histones can load onto nucleosomes both directly and through first interacting with linker DNA followed by sliding onto the nucleosome (**Fig. 6**). However, the second pathway requires a histone chaperone.

Previous single-molecule total internal reflection fluorescence (smTRIF) studies have provided insight into H1.0 interactions with DNA and nucleosomes. A smTIRF study showed that H1.0 remained bound to DNA and nucleosomes for minutes ^46^. In addition, Nap1 facilitated dissociation from DNA at super-stoichiometric concentrations of H1.0 on nucleosomes ^46^. This study is consistent with our findings. A separate single-molecule force-fluorescence study investigated the dynamic coalescence of H1.4 along single stranded-DNA that was induced by force. They find that H1.4 localizes at the junction between duplex and single stranded ^50^, which could be important for linker histone facilitated genome condensate formation. Our findings are not mutually exclusive with this complementary study. Instead, there are likely multiple mechanisms by which linker histones interact with chromatin and regulate genome accessibility.

*In vivo*, nucleosomes are spaced with linker DNA lengths of 30 to 60 bp ^51^. However, our study investigated H1.0 interactions with nucleosomes that are spaced by kilobases of DNA so the dynamics of individual linker histones and nucleosomes could be visualized and quantified. The increased density of nucleosomes will likely increase the probability of direct loading onto nucleosomes while decreasing the distance along the DNA that H1.0 must diffuse to encounter a nucleosome. We, therefore, anticipate that H1.0 loading within chromatin *in vivo* will be more efficient through both nucleosome loading models (**Fig. 6**).

The findings here provide a foundation for investigating the role of linker histones in regulating genome accessibility. For example, humans have 11 linker histone isoforms ^13^. Determining the dynamics of the different linker histones with DNA and chromatin, and then pathways used for loading onto nucleosome will be important for understanding how different linker histone isoforms are targeted to chromatin and differentially regulate genome accessibility. In addition to Nap1, there are multiple reported linker histone chaperones, including prothymosin-α ^31,52^ and sNASP ^53^. Prothymosin-α has been reported to regulate H1.0 binding to nucleosomes ^24^. It will be important to investigate the impact of these other histone chaperones on the pathways for H1.0 loading onto nucleosomes and if they are distinct from Nap1. Finally, both linker and core histones are significantly post-translationally modified (PTM), which is important for genome function and disease. Building off these findings to determine the influence of histone PTMs on linker histone dynamics and loading pathways is an important future research direction.

## METHODS

### H1 Expression and Purification

H1.0 was recombinantly expressed as previously described (**Fig. S7A**) ^3^. Briefly, for labeling, a Serine in the N-Terminal domain was mutated to a Cystine (S22C) using site-directed mutagenesis. BL21(DE3) pLysS competent E-Coli were transformed with the H1.0 expression vector that includes the S22C mutation. Single colonies of transformed E-Coli were selected on an agar plate inoculated with ampicillin. Individual transformed colonies were grown in LB growth media to an OD_600_ of 0.4-0.6 at 37C. Expression was induced by adding 0.4mM IPTG and incubation for three hours at 37C. Successful expression was confirmed using 16% acrylamide sodium dodecyl sulfate (SDS) gel electrophoresis. Following expression, the cells were pelleted at 4000g for 10 minutes, the pellet was flash frozen in liquid nitrogen, and stored in -80C for less than a week prior to purification.

To purify the expressed H1, the cell pellets were thawed on ice and resuspended in H1 lysis buffer (50mM Tris-HCl pH 8, 500mM NaCl, 10% Glycerol, 1mM EDTA, 5mM 2-Mercaptoethanol, 0.5mM PMSF). Before lysis, a complete protease inhibitor cocktail tablet (Roche) was added to 50mL of lysis buffer. The cells were lysed by sonication with a micro-needle, and the cell lysate was pelleted at 23000g for 20min at 4C. The supernatant containing soluble H1 was purified using Bio-Rex 70, 50-100 mesh, cation exchange resin packed into an FPLC-grade column. After stabilizing the resin with lysis buffer, the supernatant was added to bind H1.0 (S22C) to the column, which was then eluted off the column with a NaCl gradient using elution buffer (50mM Tris-HCl pH 8, 1M NaCl, 10% Glycerol, 1mM EDTA, 5mM 2-Mercaptoethanol, 0.5mM PMSF).

The eluted fractions were analyzed on a 16% acrylamide SDS gel, and the pooled fractions were diluted with dilution buffer (50mM Tris-HCl pH 8, 0mM NaCl, 10% Glycerol, 1mM EDTA, 5mM 2-Mercaptoethanol, and 0.5mM PMSF) so that the NaCl concentration would be reduced to ∼500mM for a second round of chromatography purification using Bio-Rex 70, 100-200 mesh (Bio-Rad), cation exchange resin. The second round of purification was performed using the same protocol used in the first round. The eluted fractions were analyzed on a 16% acrylamide SDS gel, and the pooled fractions were combined and dialyzed into MilliQ water with 2mM 2-Mercaptoethanol and 0.5mM PMSF in 4°C overnight and lyophilized. The lyophilized H1 mutants were resuspended in the storage buffer (20mM Sodium Phosphate pH7, 300mM NaCl, and 1mM EDTA), and individual aliquots were flash frozen with 10% glycerol for storage in -80°C.

### H1.0 Labeling

Each H1 mutant was labeled with Cy3 maleimide as previously described ^37^. Briefly, purified H1.0(S22C) was incubated with 10mM TCEP to ensure the mutated Cystine was reduced before the labeling reaction. Next, H1.0(S22C) was dialyzed overnight into labeling buffer (5mM PIPES pH6.1 and 2M NaCl), and mixed with 1.5M HEPES pH7.1 buffer following purging with Argon for 20min to remove the oxygen from the reaction solution. The HEPES buffer was added to the unlabeled H1.0(S22C) at a final concentration of 100mM immediately before adding 20mM Cy3-maleimide dissolved in DMF at a 7-excess molar concentration. The labeling reaction was incubated at room temperature for 1h, before incubation overnight at 4°C. The following day, the reaction solution was diluted with the binding buffer (50mM Tris-HCl pH 8, 250mM NaCl, 10% Glycerol, 1mM EDTA, 5mM 2-Mercaptoethanol, and 0.5mM PMSF) and mixed with the Bio-Rex 70, 50-100 mesh (Bio-Rad), cation exchange resin to enable binding of the labeled H1.0(S22C). The excess dye was washed with the binding buffer buffer, following by eluting the labeled H1 with elusion buffer (50mM Tris-HCl pH 8, 1M NaCl, 10% Glycerol, 1mM EDTA, 5mM 2-Mercaptoethanol, and 0.5mM PMSF). Fractions were analyzed on a 16% acrylamide SDS gel, and the pooled fractions were dialyzed into MilliQ water containing 2mM 2-Mercaptoethanol and 0.5mM PMSF, and lyophilized. Lyophilized Cy3-labeled H1.0 was resuspended in the storage buffer and individual aliquots were flash frozen with 10% glycerol for storage in -80°C (**Fig. S7A**).

### H1.0 Concentration Determination

H1.0 concentration was determined using the modified Lowry assay based on H1.0 samples with known concentrations between 100ng/uL to 1000ng/uL. The estimated concentrations were verified by examining equal amounts of purified H1.0 and the reference H1.0 on a 16% acrylamide SDS gel.

### Histone Octamer Preparation and Labeling

Human H2A(K119C), H2B, H3(C110A), and H4 were purchased from the Histone Source at Colorado State University. The lyophilized histones were resuspended in unfolding buffer (20mM Tris-HCl pH7.4, 7M Guanidinium, 1mM DTT) at 5mg/mL, and the unfolded histones were mixed at 1:1.2 mass ratio between H3:H4 tetramer and H2A(K119C):H2B heterodimer. Successful refolding of octamers was achieved by an overnight double dialysis into octamer refolding buffer (20 mM Tris-HCl pH 7.4, 1 mM EDTA, 2 M NaCl, and 5 mM 2-Mercaptoethanol). The refolded octamer was recovered and, H2A(K119C) was labeled with Cy5 maleimide using the same protocol described above for H1.0(S22C) labeling with Cy3. Prior to purifying the labeled octamer from the heterodimer and free histones, and labeling reaction was quenched with 10mM DTT. The quenched reaction was purified using gel-filtration FPLC chromatography with a Superdex^TM^ 200 Increase 10/300 GL gel-filtration column (GE Healthcare), and the fractions were analyzed on a 16% acrylamide SDS gel. The pooled fractions were buffer exchanged and concentrated in octamer storage buffer 0.5X TE (10mM Tris-HCl pH 8.0, 1mM EDTA), and stored in -20°C with 40% glycerol (**Fig. S7B**).

### Nap1 Expression and Purification

Rosetta2 (DE3) competent *E. coli* were transformed with a mouse Nap1 expression vector (a gift from Stefan Dimitrov). Single colonies were selected on an agar plate inoculated with ampicillin and chloramphenicol. Individual transformed colonies were grown in LB growth media to an OD_600_ of 0.4-0.6 at 37°C. Expression was induced by adding 0.25mM IPTG and incubation for three hours at 37°C. Successful expression was confirmed using 12% acrylamide sodium dodecyl sulfate (SDS) gel electrophoresis. Following expression, the cells were pelleted at 4000g for 10 minutes, the pellet was flash frozen in liquid nitrogen, and stored at -80°C for less than a week before purification.

To purify the expressed Nap1, the cell pellets were thawed on ice and resuspended in Nap1 buffer A (20mM Tris-HCl pH 7.4, 20mM imidazole, 100mM NaCl, 10% Glycerol, 1mM BME, 0.5mM PMSF). Before lysis, two complete Roche EDTA-free protease inhibitor cocktail tablets were added to 30mL of Nap1 buffer A. The cells were lysed by sonication with a micro-needle, and the cell lysate was pelleted at 23000g for 15min at 4°C. The supernatant containing soluble Nap1 was purified using a Nickel column (Cytiva). After stabilizing the column with Nap1 buffer A, the supernatant was added to bind to the column, which was then eluted off the column with Nap1 buffer B (20mM Tris-HCl pH 7.4, 500mM imidazole, 100mM NaCl, 10% Glycerol, 1mM BME, 0.5mM PMSF).

The eluted fractions were analyzed on a 12% acrylamide SDS gel. The pooled fractions were further purified using ion exchange chromatography with a MonoQ column (Cytiva). The pooled Nap1 fractions were bound to the column in Nap1 buffer C (20mM Tris-HCl pH 7.4, 100mM NaCl, 10% Glycerol, 1mM BME, 0.5mM PMSF) and eluted with Nap1 buffer D (20mM Tris-HCl pH 7.4, 1M NaCl, 10% Glycerol, 1mM BME, 0.5mM PMSF). The eluted fractions were analyzed on a 12% acrylamide SDS gel, and the pooled fractions were combined and concentrated in Nap1 buffer C. Iindividual aliquots were flash frozen with 10% glycerol for storage in -80°C (**Fig. S7C**).

### DNA Preparation and Purification

All short DNA constructs were synthesized using polymeric chain reaction with Pfu polymerase. To prepare the 147 bp carrier DNA for lambda DNA chromatinization, unmodified forward (5′-GCATAATTCTCTTACTGTCATGCCATCCG-3′) and reverse (5′-CTGCTATGTGGCGCGGTATTATCC-3′) primers (IDT) were used to amplify from pUC19 plasmid as the template. A second carrier DNA (179bp in length) was amplified for lambda DNA chromatinization using unmodified forward (5′-CTGTGCCACCACGGAACAGGATGTATATATCTGACACGTGCCTGG-3′) and reverse (5′-GAAGAGGTGGCGCGTAACTGGAGAATCCCGGTGCCG-3′) primers (IDT) and a pUC19 plasmic containing Widom 601 DNA ^54^ as the template. All PCR reactions were purified using anion exchange chromatography with a Gen-Pak^TM^ Fax 4.6×100mm column (Waters) and buffer exchanged into the 0.5X TE storage buffer using a 30kDa Amicon filter (Millipore).

### Lambda Tether Preparation

To synthesize lambda tether with multi-biotin modification on each end, the 12-base 5′ single-stranded overhangs on each end of the bacteriophage λ DNA were filled with the DNA polymerase I Klenow fragment that lacks 5′→3′ endonuclease activity. The reaction solution contained 6nM Lambda DNA, 100µM dATP, 100µM dTTP, 100µM dGTP, 80µM biotin-14-dCTP and 0.075 U/µL Klenow fragment in 1X NEB2 buffer to enable incorporation of biotinylated nucleotides during the polymerase reaction. The reaction solution was incubated at 37°C for 30min followed by heat inactivation at 75°C for 20min. Following the filling of the single-stranded lambda DNA ends, the enzyme was removed using phenol extraction followed by exchanging into 0.5X TE at 4°C. The concentrated tether was aliquoted and stored at -20°C.

### Nucleosome Preparation

NaCl gradient dialysis was used to reconstitute nucleosomes along the prepared lambda tethers with multi-biotin ends as previously implemented ^55^. The chromatinization density was regulated by changing the concentrations of the 147bp low-affinity carrier DNA, the 187bp 601 carrier DNA, and the Cy5-Octamer in the reconstitution mixture. To include sparse numbers of nucleosomes along the Lambda tether, the 187bp 601 carrier DNA was included in the reconstitution mixture to control the number nucleosome reconstituted into the lambda DNA. Lambda tether, carrier DNA and Cy5-octamer mixture in refolding buffer (10mM Tris-HCl pH 8.0, 1mM EDTA, 2M NaCl, and 1mM Benzamidine-HCl) were reconstituted into the buffer condition without NaCl (10mM Tris-HCl pH 8.0, 1mM EDTA, and 1mM Benzamidine-HCl) at 4°C for 4-6 hours using double dialysis. The dialysis bag was moved to a fresh buffer condition without NaCl to continue the NaCl gradient dialysis overnight at 4°C. The reconstituted nucleosomes were analyzed by agarose gel electrophoresis to confirm the formation of Cy5-labeled nucleosomes along the lambda DNA tether (**Fig. S8**).

### Dual optical tweezers and fluorescence confocal imaging experimental system

Single-molecule experiments were performed using a dual optical trap C-Trap instrument (LUMICKS). A microfluidic flow cell was implemented to separately introduce each reagent needed for a single molecule test. Low Reynolds Number laminar flow enabled minimal mixing between different reagents within the flow cell, while a computer-controlled stage enabled rapid movement of the two optical traps within the flow cell. Streptavidin-coated polystyrene beads (4.34µm in size, Spherotech), biotinylated lambda tethers, and imaging buffer (50mM HEPES pH 7.4, 130mM NaCl, 0.005% Tween-20, 2mM Trolox, 450 ug/mL glucose oxidase, 22ug/mL catalase, 0.0115% v/v COT (Cyclooctatetraen), 0.012% v/v NBA (3-Nitrobenzyl alcohol), 1.6% w/v glucose) were introduced in parallel in channels 1 through 3, respectively. An additional, orthogonal channel 4 was used to introduce 3nM Cy3-H1.0 diluted in the imaging buffer. Laminar flow was introduced while the traps were held in channel 1 to enable trapping of a single streptavidin-coated bead in each optical trap. To calibrate the trap stiffness, the beads were moved to channel 3 while the flow was stopped. A spectral analysis was performed to achieve trap stiffness of ∼0.2-0.4pN/nm for each trap. The stage was moved to bring the trapped beads to channel 2 to enable the successful capture of a single lambda tether between the two streptavidin-coated beads. Binding interactions between the biotinylated dCTPs on the tether ends and streptavidin on the surface of the beads enabled stable capturing of the lambda tether between the two beads. The presence of a captured lambda tether was detected by probing the tether force-extension curve as the two beads were subjected to repeated approach-retraction movement with respect to each other. The flow was stopped, and the captured tether was brought into channel 3 to verify the presence of a single tethered molecule using a lambda DNA worm-like chain reference model. For experiments with nucleosome-containing tethers, confocal line scanning along the tether was implemented using the 640nm laser to visualize the location of the reconstituted nucleosomes. The mechanical force corresponding to each test was set using the Bluelake software (LUMICKs). Before transferring the tether to the H1.0-Cy3 channel, the 640nm excitation laser was turned off, so only the 532nm excitation laser (green) remained on. This allowed for the H1.0-Cy3 molecules to be visualized following the transfer of the tether into the H1.0-Cy3 channel. The stage was then moved to bring the assembled construct into the orthogonal channel 4 containing Cy3-H1 to monitor H1 binding and one-dimensional diffusion.

### H1.0 single-molecule tracking and data analysis

HDF5 format files were processed using the commercial Lakeview software (LUMICKs) along with the commercial Pylake Python package (LUMICKs) to detect individual traces. For each individual H1.0-Cy3 tracked trace, the extracted position and time data were further processed to quantify the dwell time and diffusion coefficient using a custom-built MATLAB code.

The mean-square displacement (MSD) for each tracked particle was calculated using:

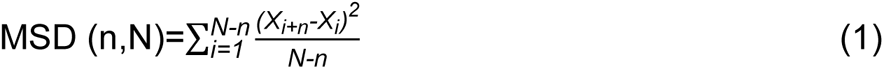

where N is the total number of time points for each trace, *n* is the MSD time-lag size (n=1→N-1), *i* is the window of displacement measurement, and *X_i_* is the estimated position of each tracked H1.0-Cy3 in the frame *i*. Assuming H1.0 exhibits Brownian diffusion, the diffusion coefficient (D) for each tracked H1.0 was estimated using an ordinary least square estimator by solving for D in the following equation, MSD=2Dt, based on MSD measured within the first 1s window as previously described ^56^.

### Position measurements of immobile DNA-bound proteins

Cy5 labeled histone octamer was reconstituted onto lambda DNA to provide a reference for immobile DNA-bound protein. Position of Cy5-labeled nucleosomes was quantified when the tether was under 1pN, 3pN and 10pN of force to represent immobile tether-bound molecules. Time traces (n = 354) of Cy5-labeled nucleosomes were quantified (146 under 1pN, 151 under 3pN, and 57 under 10pN). The estimated mean-squared displacement of a single molecule over time t (MSD(t)) can be described as:

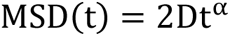

where D is the diffusion coefficient and α is the temporal scaling exponent ^57^. The MSD was determined with a time step of 40ms over a time window of 1s. For a molecule exhibiting unconstrained Brownian motion, α = 1, while for a completely immobile molecule, α = zero. We fit the averaged MSD vs. time to the above equation to determine α for each nucleosome. The values of α and the MSD at t = 1s (MSD_t=1s_) for each nucleosome under either 1pN, 3pN, and 10pN were indicated on a scatter plot (**Figure S1B**). The maximum recorded value of α for a single nucleosome (0.22, 0.22 and 0.20 under 1pN, 3pN and 10pN, respectively) was used as the cut-off threshold for determining whether a tracked H1.0 is immobile (**Figure S1C**, shown for 1pN only). There was a small fraction (less than 5%) of H1.0 traces that had an α of less than the cut-off threshold but an MSD_t=1s_ that was larger than the maximum MSD_t=1s_ for nucleosomes. Upon visual examination, these traces showed incorrectly tracked positions and were therefore not included (**Figure S1C**, black squares, shown for 1pN only).

### Statistical analysis

In order to report uncertainties associated with each fit parameter, each data set was randomly divided into three subgroups with similar sample size based on selection without replacement statistical analysis. All reported uncertainties are in the mean ± SEM (standard error of the mean) format. The reported *p*-values are based on student’s t-test performed in MATLAB.

## DATA AVAILABILITY

The raw data files pertaining to the data presented in the main and supplementary figures in this manuscript can be made available upon request from the authors.

## ACKNOWLEDGEMENTS

The authors acknowledge the feedback and insights provided by the Poirier Lab members and the members of the LUMICKS team. We are grateful to Stefan Dimitrov for the mouse Nap1 expression vector. We would like to acknowledge the staff at the nano systems laboratory at the Ohio State University Department of Physics. Partial funding for shared facilities used in this research was provided by the Center for Emergent Materials: an NSF MRSEC under award number DMR-2011876. Support for the LUMICKS C-Trap was provided by The National Institutes of Health (S10OD028705 to M.G.P.). This work was funded by the National Institutes of Health (R35 GM139564 to M.G.P.); National Science Foundation (MCB 2411725 to M.G.P.); Funding for open access charge: National Institutes of Health R35 GM139654.

## SUPPLEMENTARY INFORMATION

### Supplementary Figures

**Figure S1.**
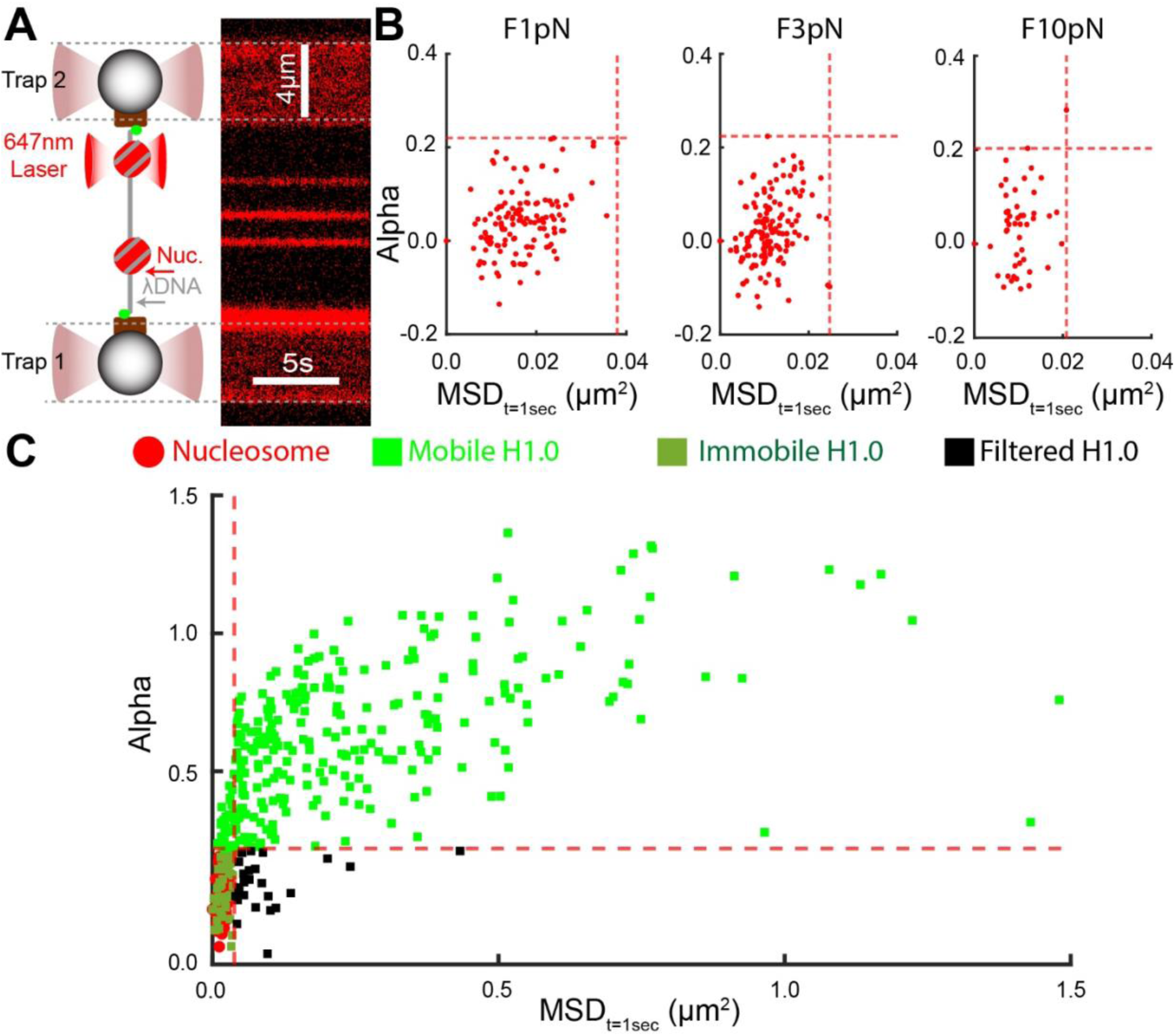
Nucleosome position calibration measurements for differentiating between mobile and immobile H1.0. (**A**) Illustration of the single-molecule assembly along with a kymograph of Cy5-labled nucleosomes sparsely reconstituted along the nucleosome-containing tether. **B)** Scatter plots of MSD_t=1sec_ (mean squared displacement at 1 second) versus α for nucleosomes while the tether was subjected to 1pN, 3pN, or 10pN mechanical force. **C)** Scatter plot of MSD_t=1sec_ versus α for H1.0 (dark green, bright green and black) on DNA-only tethers and the nucleosomes (red) within nucleosome containing tethers that are subject to a constant force of 1pN. The tracked H1.0 traces were characterized as mobile (bright green), immobile (dark green), or filtered (falsely detected position, black).

**Figure S2.**
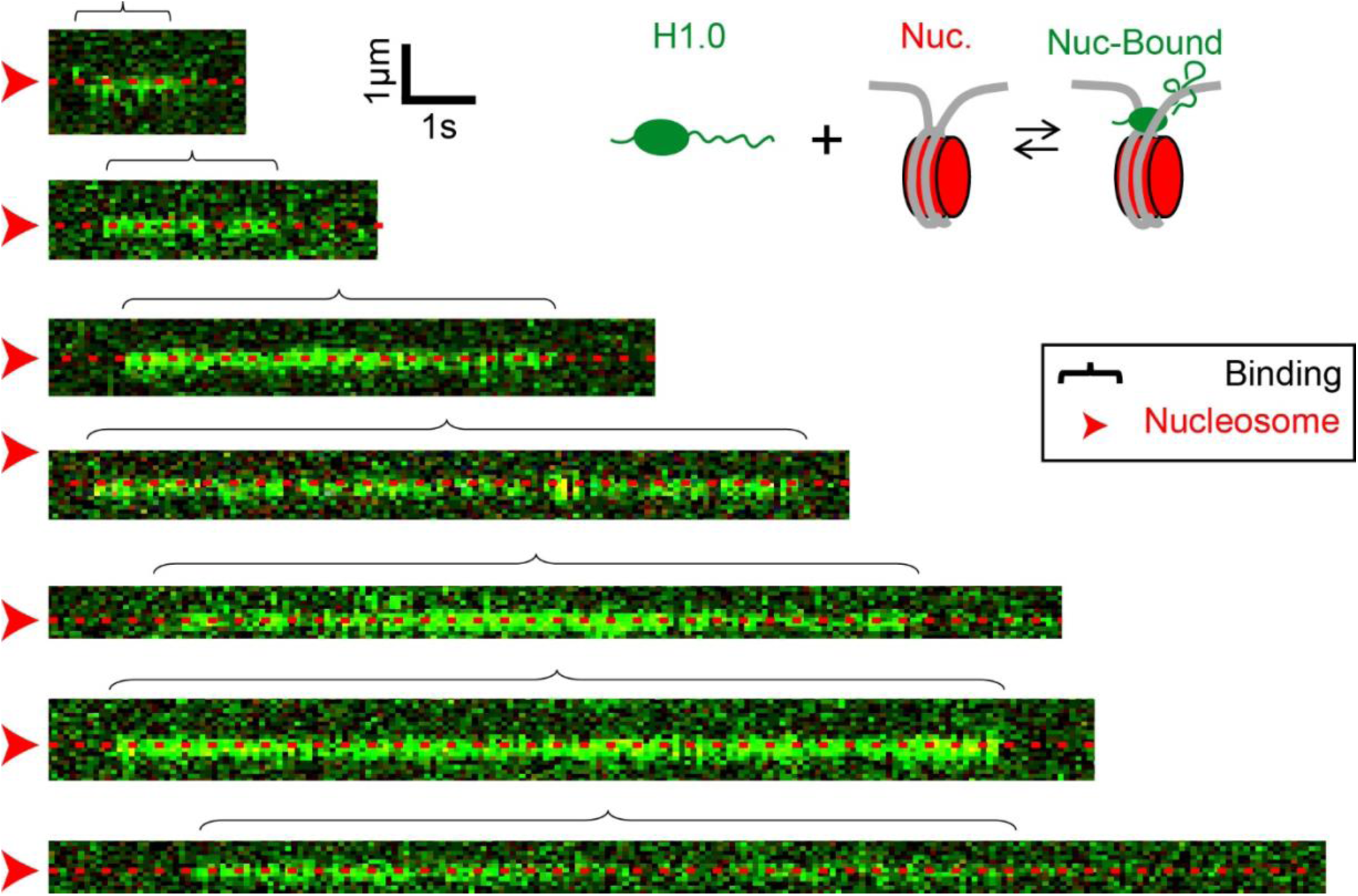
Representative kymographs of H1.0 that directly load onto nucleosomes when first binding a nucleosome-containing tether. The nucleosome positions are indicated with the red arrowhead along with the red dash lines. The illustration provides a visual description of the observed binding events. H1.0 is immobile upon direct binding to a nucleosome. All measurements were done while the tether was under a constant force of 1pN.

**Figure S3.**
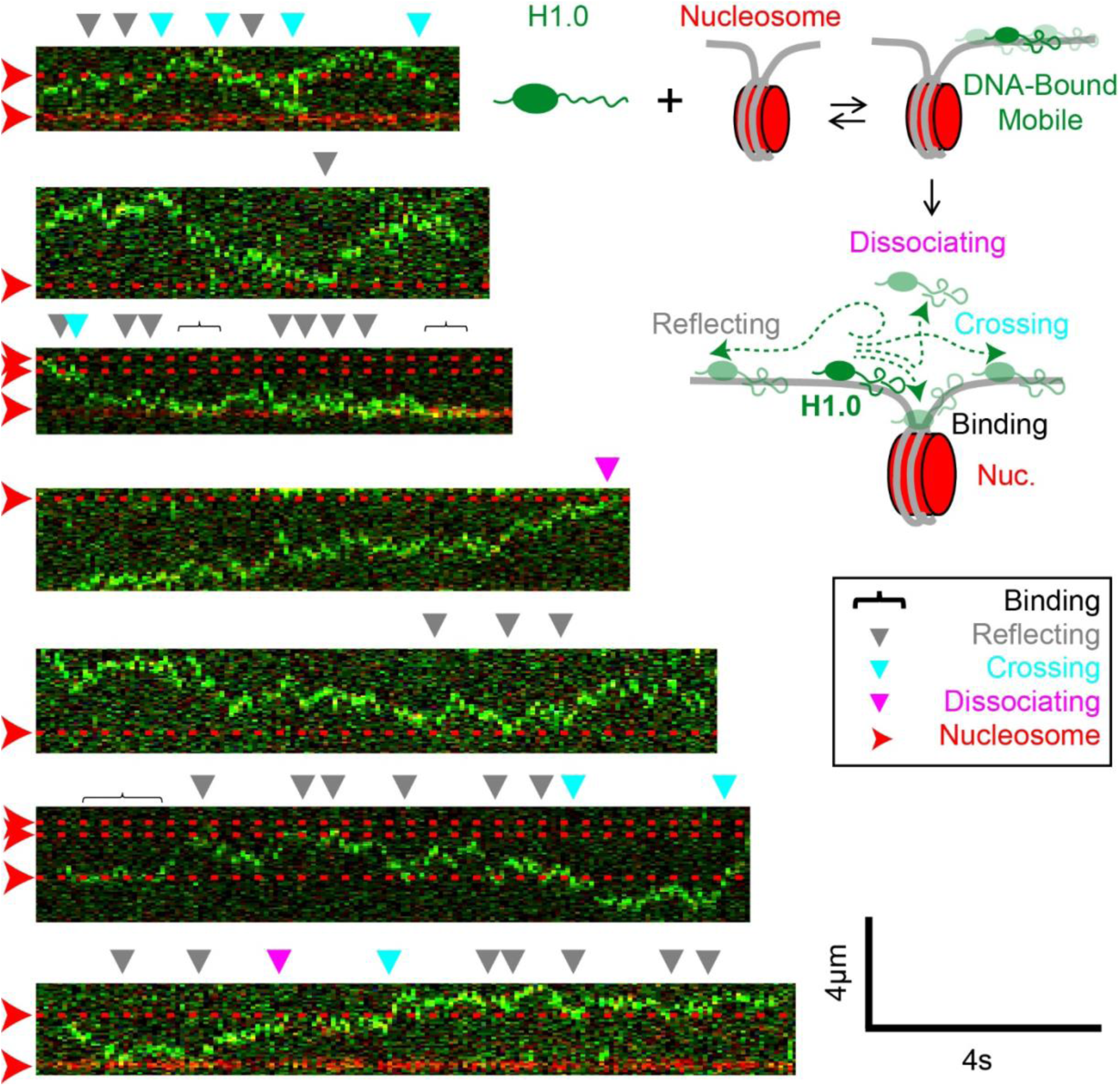
Representative kymographs of H1.0_mobile_ that directly load onto DNA when first binding a nucleosome-containing tether. The nucleosome positions are indicated with the red arrowhead along with the red dash lines. The illustration provides a visual description of the observed binding events. Following H1.0mobile colocalization with a nucleosome, H1.0mobile can bind to (black bracket), reflect off (grey arrow), cross over (cyan arrow), or dissociate from (magenta arrow) the nucleosome. H1.0_mobile_ reflects off nucleosomes more often than binding to, crossing over or dissociating from nucleosomes. All measurements were done while the tether was under a constant force of 1pN.

**Figure S4.**
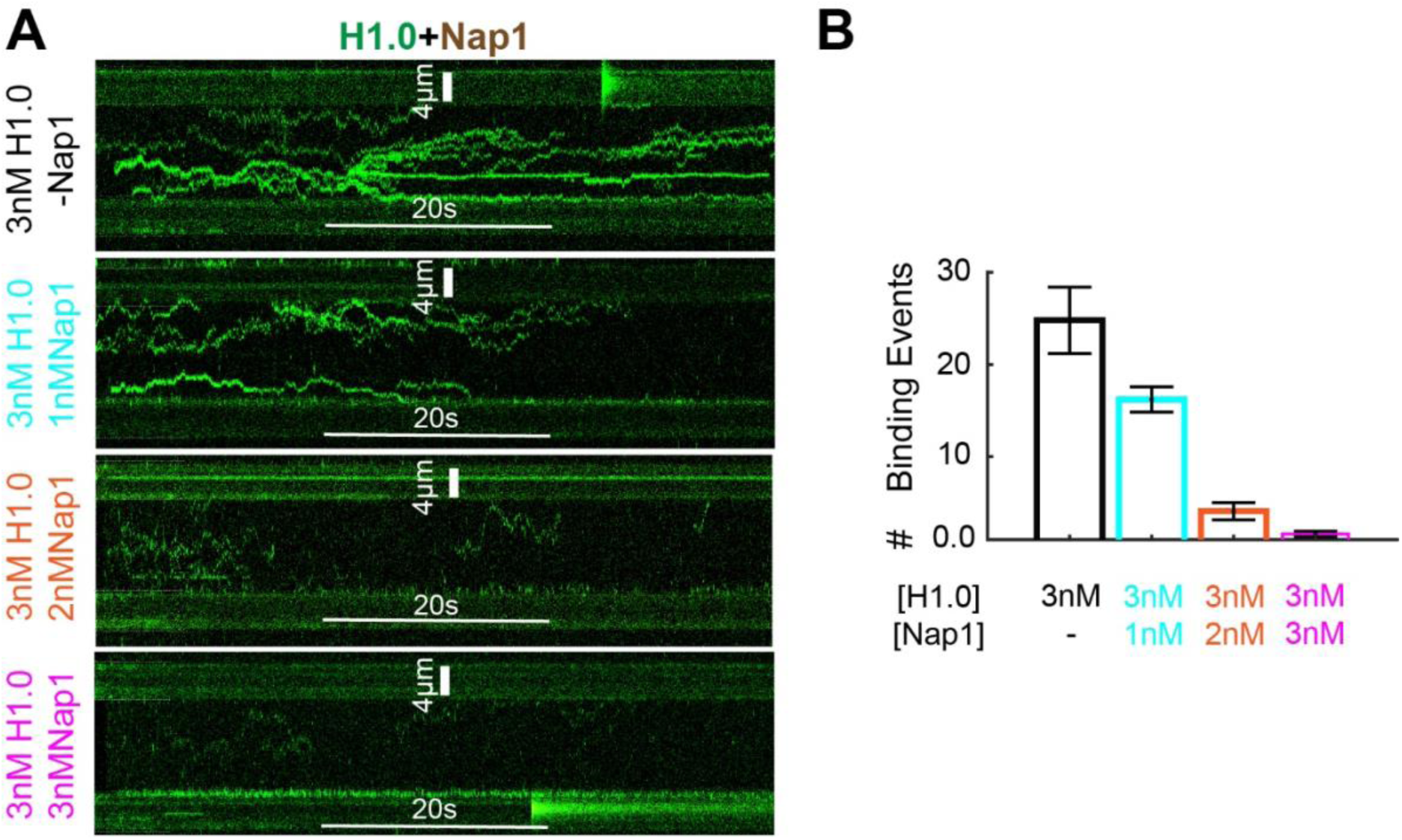
Elevated NAP1 concentration prevents binding interaction between H1.0 and dsDNA. (**A**) Representative kymographs of H1.0 that directly load onto DNA-only tethers at Nap1 concentrations of 0nM (black), 1nM (cyan), 2nM (orange), and 3 nM (magenta). Elevated NAP1 concentration reduces H1.0 occupancy on the DNA-only tether. (**B**) Number of binding events to the DNA-only tether over a period of 3min. The number of H1.0 binding events to the DNA-only tether systematically reduced with increased NAP1 concentration. All measurements were done while the tether was under a constant force of 1pN.

**Figure S5.**
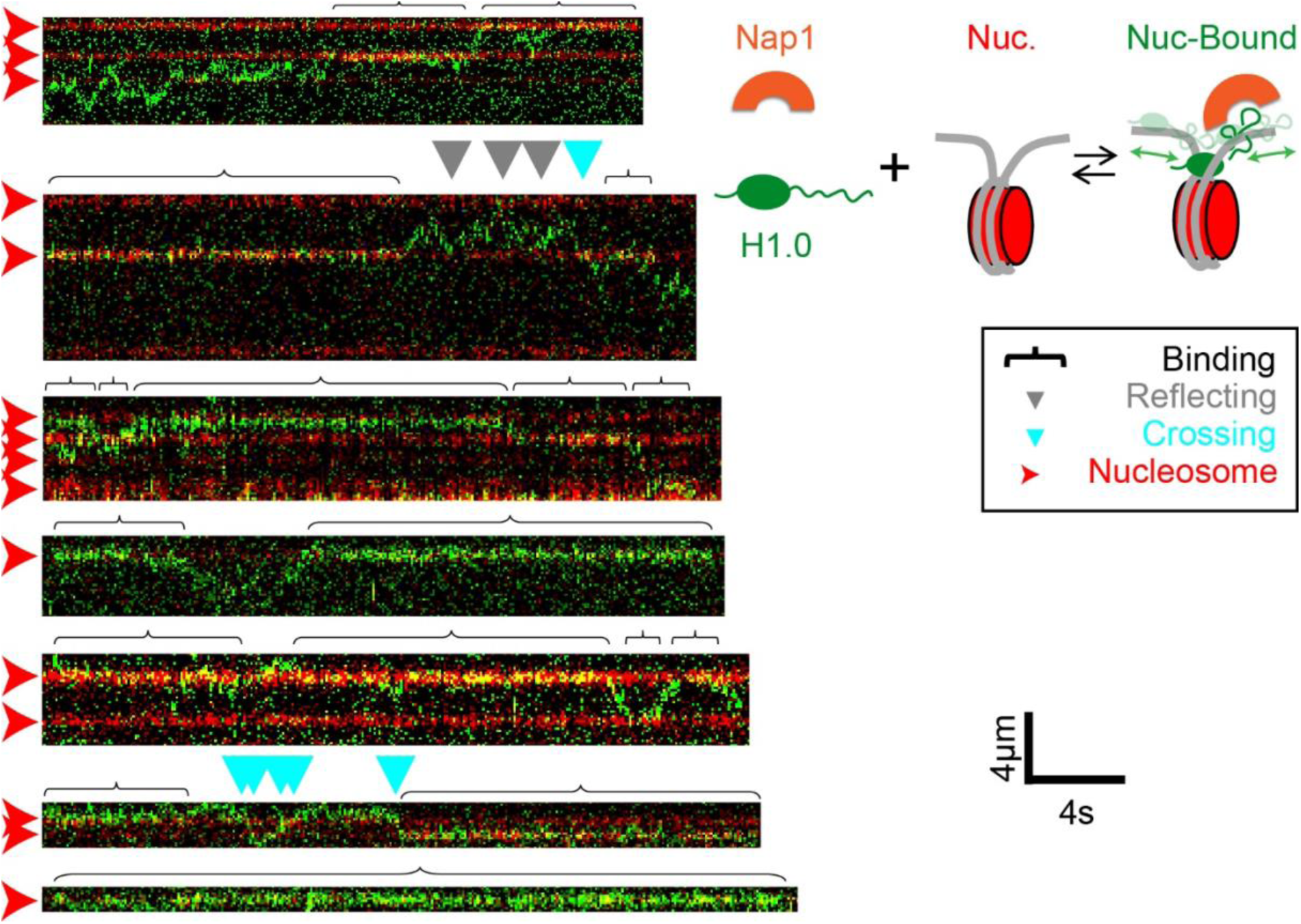
Representative kymographs of H1.0 that directly load onto nucleosomes when first binding a nucleosome-containing tether in the presence of Nap1. The nucleosome positions are indicated with the red arrowhead along with the red dash lines. The illustration provides a visual description of the observed binding events. Following H1.0_mobile_ colocalization with a nucleosome, H1.0_mobile_ is observed to bind to (black bracket), reflect off (grey arrow), or cross over (cyan arrow) the nucleosome. H1.0 remains bound to the nucleosome for many second but also transitions to DNA and then either rebinds the same nucleosome or a neighboring nucleosome. All measurements were done while the tether was under a constant force of 1pN and in the presence of 2nM Nap1.

**Figure S6.**
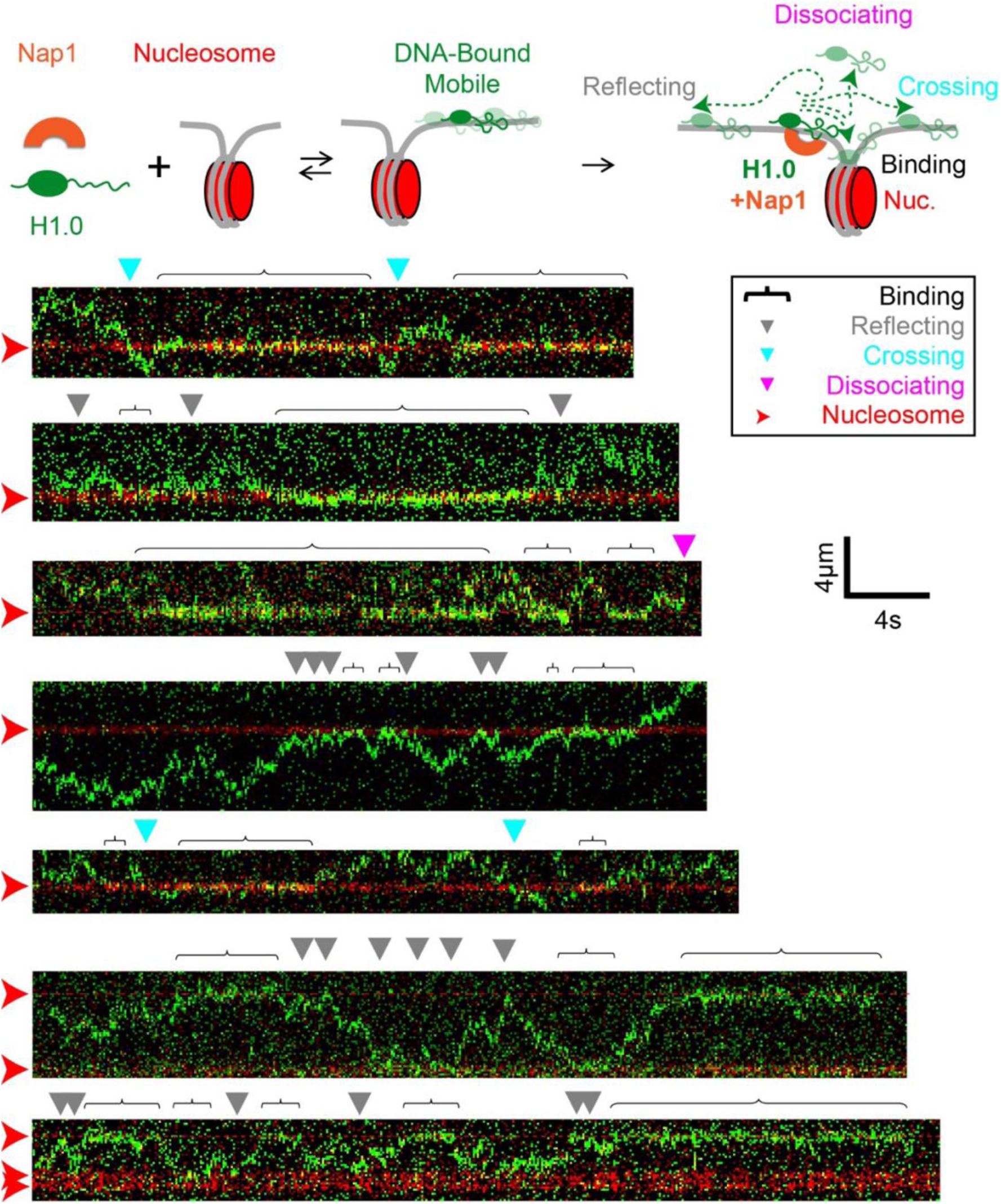
Representative kymographs of H1.0_mobile_ that directly load onto DNA when first binding a nucleosome-containing tether in the presence of Nap1. The nucleosome positions are indicated with the red arrowhead along with the red dash lines. The illustration provides a visual description of the observed binding events and the outcomes following an H1.0_mobile_-nucleosome colocalization event. Following H1.0_mobile_ colocalization with a nucleosome, H1.0_mobile_ can bind to (black bracket), reflect off (grey arrow), cross over (cyan arrow), or dissociate from (magenta arrow) the nucleosome. In the presence of Nap1, H1.0_mobile_ largely binds to or reflects off nucleosomes. All measurements were done while the tether was under a constant force of 1pN.

**Figure S7.**
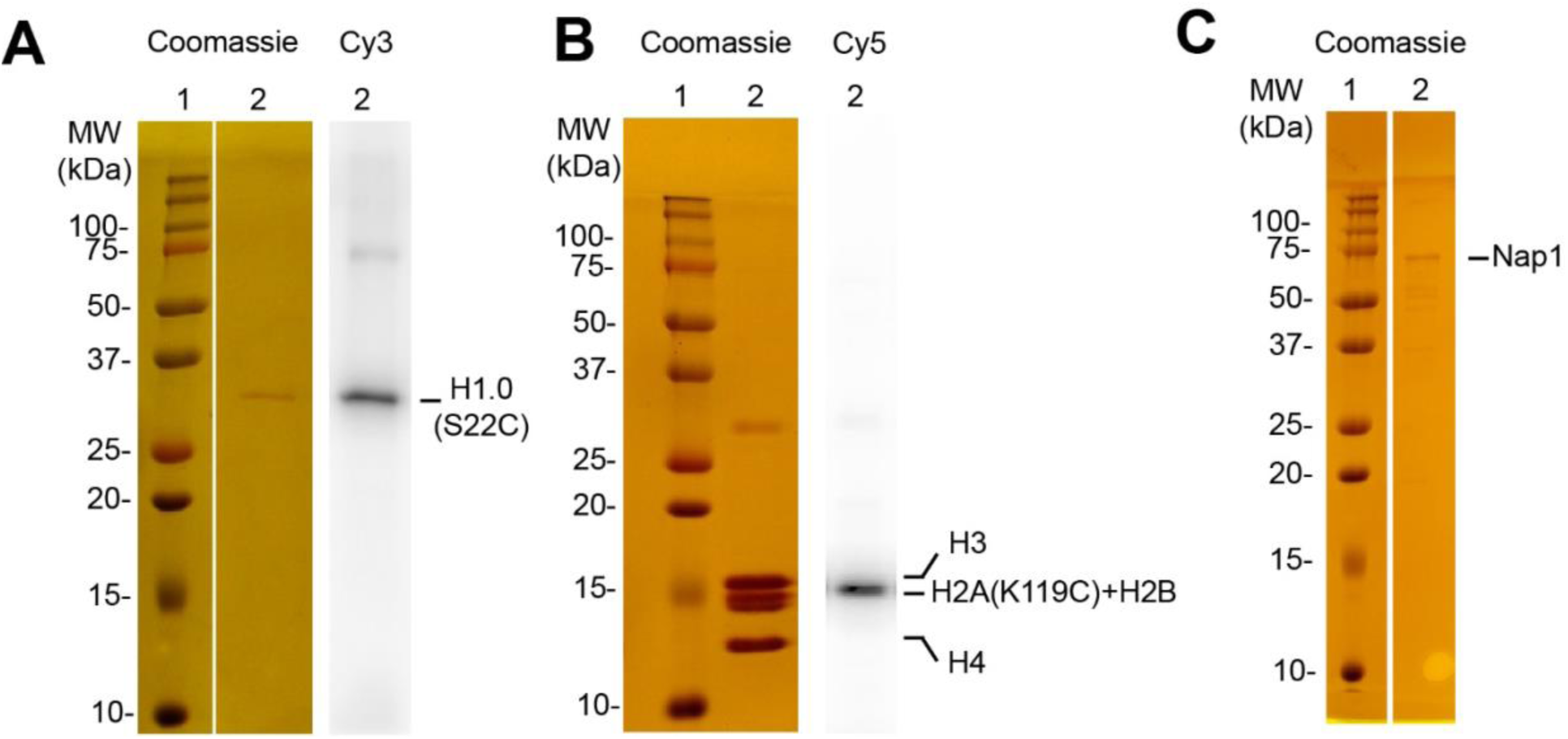
Purified H1.0(S22C)-Cy3, histone octamer-Cy5, and Nap1. **(A)** SDS-PAGE analysis of H1.0(S22C) by Coomassie Brilliant Blue imaging. Lane 1: standard protein molecular weights in kDa. Lane 2: Purified human H1.0 visualized on SDS acrylamide gel. The Cy3 labeling of H1.0(S22C) was confirmed by Cy3 fluorescence (Lane 2, Cy3). **(B)** SDS-PAGE analysis of the histone octamer by Coomassie Brilliant Blue imaging. Lane 1: standard protein molecular weights in kDa. Lane 2: Purified human histone octamer visualized on SDS acrylamide gel. The Cy5 labeling of H2A(K119C) was confirmed by Cy5 fluorescence (Lane 2, Cy5). **(C)** SDS-PAGE analysis of Nap1 by Coomassie Brilliant Blue imaging. Lane 1: standard protein molecular weights in kDa. Lane 2: Purified mouse Nap1 visualized on SDS acrylamide gel.

**Figure S8.**
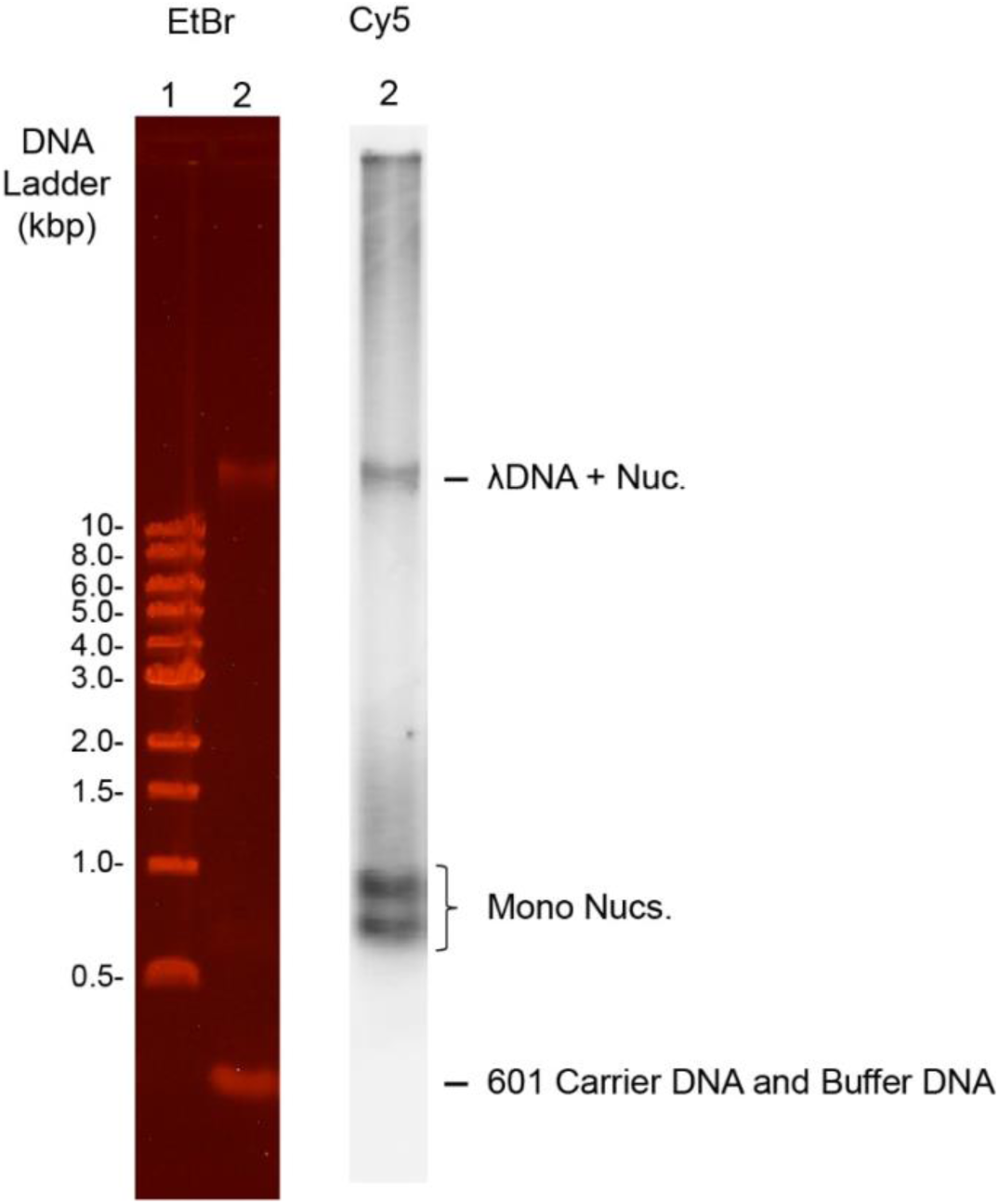
Gel electrophoresis characterization of nucleosome reconstitution along lambda tether DNA in the presence of excess 601 carrier DNA and low-affinity carrier DNA. Agarose gel electrophoresis analysis with ethidium bromide imaging of the nucleosome reconstitutions onto lambda DNA. Lane 1: Standard 1kb DNA ladder. Lane 2: Nucleosomes reconstituted along lambda DNA in the presence of excess short carrier DNA. Lambda DNA tether and excess unconverted carrier DNA were visualized using EtBr staining. Fluorescence signal from H2A(K119C)-Cy5 was used to confirm the presence of nucleosomes reconstituted along the lambda DNA tether (Lane 2, Cy5).

### Supplementary Tables

**Table S1.** Summary of mobility analysis of H1.0 bound to DNA-only and nucleosome containing tethers.

| Tether Type | Force | [Nap1] | Mobile Fraction | Alpha ($\alpha$) | Diffusion Coefficient (D) |
| --- | --- | --- | --- | --- | --- |
| DNA-only | 1pN | 0 | $0.84 \pm 0.01$ | $0.80 \pm 0.02$ | $0.281 \pm 0.002 \mu\text{m}^2/\text{s}$ |
| DNA-only | 3pN | 0 | $0.91 \pm 0.01$ | $0.79 \pm 0.02$ | $0.31 \pm 0.01 \mu\text{m}^2/\text{s}$ |
| DNA-only | 10pN | 0 | $0.91 \pm 0.01$ | $0.79 \pm 0.02$ | $0.368 \pm 0.003 \mu\text{m}^2/\text{s}$ |
| DNA-only | 1pN | 1nM | 1 | $0.68 \pm 0.02$ | $0.40 \pm 0.02 \mu\text{m}^2/\text{s}$ |
| DNA-only | 1pN | 2nM | 1 | $0.92 \pm 0.01$ | $0.60 \pm 0.06 \mu\text{m}^2/\text{s}$ |
| DNA-only | 1pN | 3nM | 1 | $1.10 \pm 0.01$ | $0.95 \pm 0.29 \mu\text{m}^2/\text{s}$ |
| Nucleosome-containing | 1pN | 0 | $0.88 \pm 0.01$ | $0.84 \pm 0.01$ | $0.344 \pm 0.02 \mu\text{m}^2/\text{s}$ |
| Nucleosome-containing | 1pN | 2nM | $0.84 \pm 0.01$ | $0.79 \pm 0.01$ | $0.490 \pm 0.006 \mu\text{m}^2/\text{s}$ |

**Table S2.** Summary of dwell-time analysis of H1.0 bound to DNA-only and nucleosome-containing tethers.

| **Tether Type** | **Force** | **[Nap1]** | **Mobility** | **$\tau_{fast}$** | **$\tau_{slow}$** | **$A_{fast}$** |
| --- | --- | --- | --- | --- | --- | --- |
| DNA-only | 1pN | 0 | Mobile | $6 \pm 1$ s | $23 \pm 2$ s | $0.55 \pm 0.09$ |
| DNA-only | 1pN | 0 | Immobile | $1.4 \pm 0.3$ s | $44 \pm 3$ s | $0.49 \pm 0.04$ |
| DNA-only | 3pN | 0 | Mobile | $4.6 \pm 0.1$ s | $40 \pm 8$ s | $0.83 \pm 0.04$ |
| DNA-only | 3pN | 0 | Immobile | $2.1 \pm 0.5$ s | $40 \pm 20$ s | $0.78 \pm 0.04$ |
| DNA-only | 10pN | 0 | Mobile | $6 \pm 1$ s | $40 \pm 6$ s | $0.71 \pm 0.05$ |
| DNA-only | 10pN | 0 | Immobile | $0.8 \pm 0.2$ s | $16 \pm 5$ s | $0.4 \pm 0.1$ |
| DNA-only | 1pN | 1nM | Mobile | $6 \pm 2$ s | $14 \pm 3$ s | $0.7 \pm 0.2$ |
| DNA-only | 1pN | 2nM | Mobile | $10.7 \pm 0.8$ s | N.A. | 1.0 |
| DNA-only | 1pN | 3nM | Mobile | $3.8 \pm 0.6$ s | N.A. | 1.0 |
| Nucleosome-Containing | 1pN | 0 | Mobile | $5.7 \pm 1.4$ s | $22 \pm 0.02$ s | $0.60 \pm 0.10$ |
| Nucleosome-Containing | 1pN | 0 | Immobile | $6.6 \pm 1.2$ s | $0.79 \pm 0.10$ s | $0.62 \pm 0.05$ |
| Nucleosome-Containing | 1pN | 2nM | Mobile | $9.1 \pm 2.0$ s | $0.20 \pm 0.04$ s | $0.70 \pm 0.2$ |
| Nucleosome-Containing | 1pN | 2nM | Immobile | $4.2 \pm 1.4$ s | $0.55 \pm 0.04$ s | $0.48 \pm 0.07$ |

**Table S3.** Summary of the probability of binding (0.4s < colocalization time), crossing, reflecting, and dissociating events for H1.0_mobile_-nucleosome colocalization events and H1.0_mobile_ colocalization events at randomly selected points along DNA-only tethers.

| **Tether Type** | **Force** | **[Nap1]** | **Binding Probability** | **Crossing Probability** | **Reflecting Probability** | **Dissociating Probability** |
| --- | --- | --- | --- | --- | --- | --- |
| DNA-only | 1pN | 0 | $0.14 \pm 0.02$ | $0.41 \pm 0.01$ | $0.40 \pm 0.01$ | $0.024 \pm 0.001$ |
| DNA-only | 1pN | 2nM | $0.0865 \pm 0.002$ | $0.53 \pm 0.03$ | $0.35 \pm 0.02$ | $0.014 \pm 0.004$ |
| Nucleosome-Containing | 1pN | 0 | $0.162 \pm 0.002$ | $0.16 \pm 0.01$ | $0.650 \pm 0.004$ | $0.014 \pm 0.004$ |
| Nucleosome-Containing | 1pN | 2nM | $0.218 \pm 0.002$ | $0.291 \pm 0.006$ | $0.467 \pm 0.004$ | $0.011 \pm 0.001$ |

---

**Extended Figure 1.**
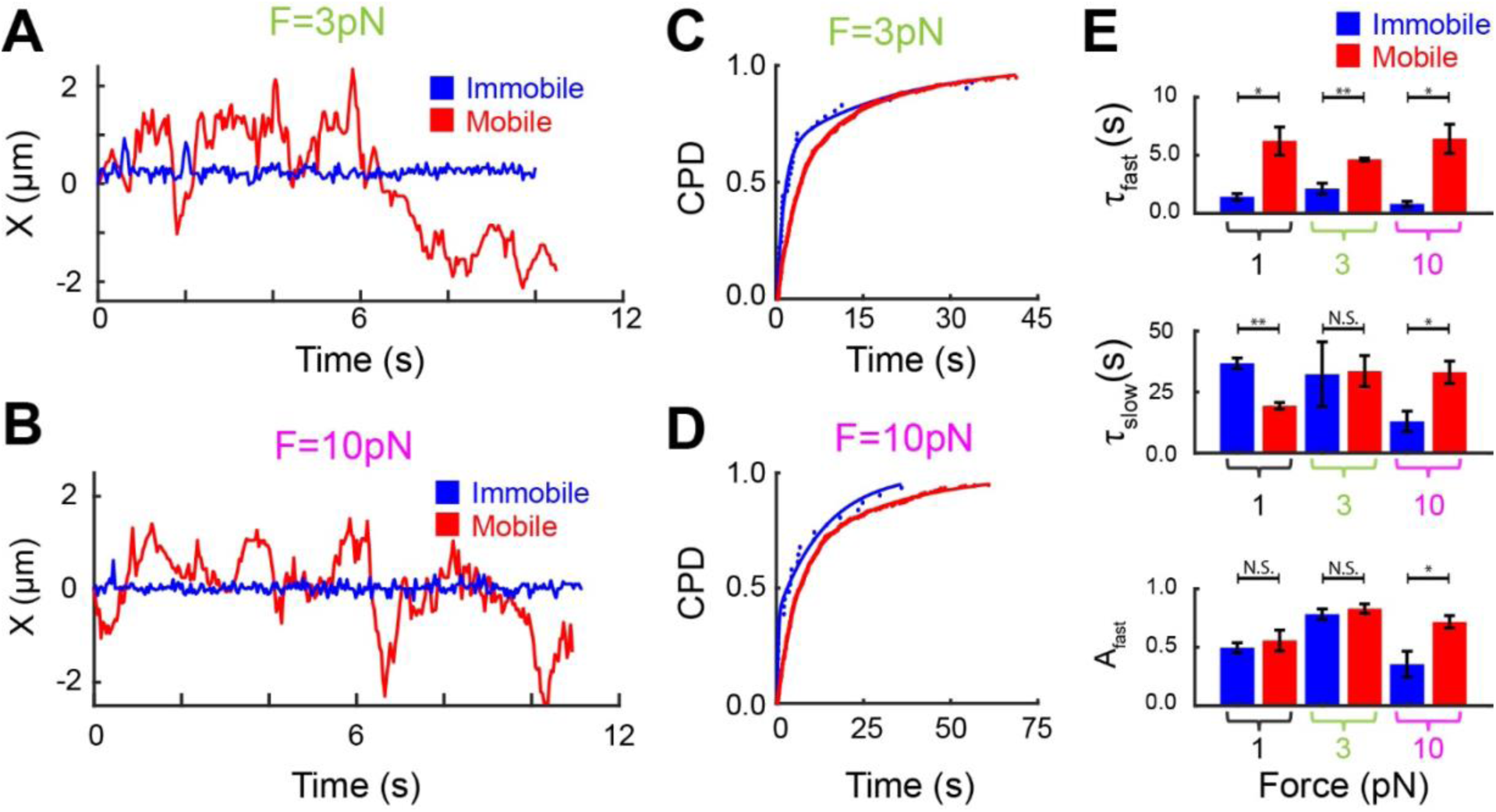
Force dependence of DNA bound H1.0 resident time . (**A**) and (**B**) Net displacement of a diffusing (red) versus immobile (blue) trace over time when bound to tether under 3pN and 10pN of constant force, respectively. (**C**) and (**D**) Dwell time CPDs for immobile (blue) versus diffusing (red) traces that were bound to DNA under 3pN and 10pN of constant force, respectively. All four dwell time CPDs were best fit to a double-exponential model to yield short dwell time (τ_fast_), long dwell time (τ_slow_) and short dwell time fraction (A_fast_). (**E)** Fit parameters τ_fast_, τ_slow_ and A_fast_ for mobile (red) versus immobile (blue) H1.0 traces under 1pN (black), 3pN (green) and 10pN (magenta) of constant force.

**Extended Figure 2.**
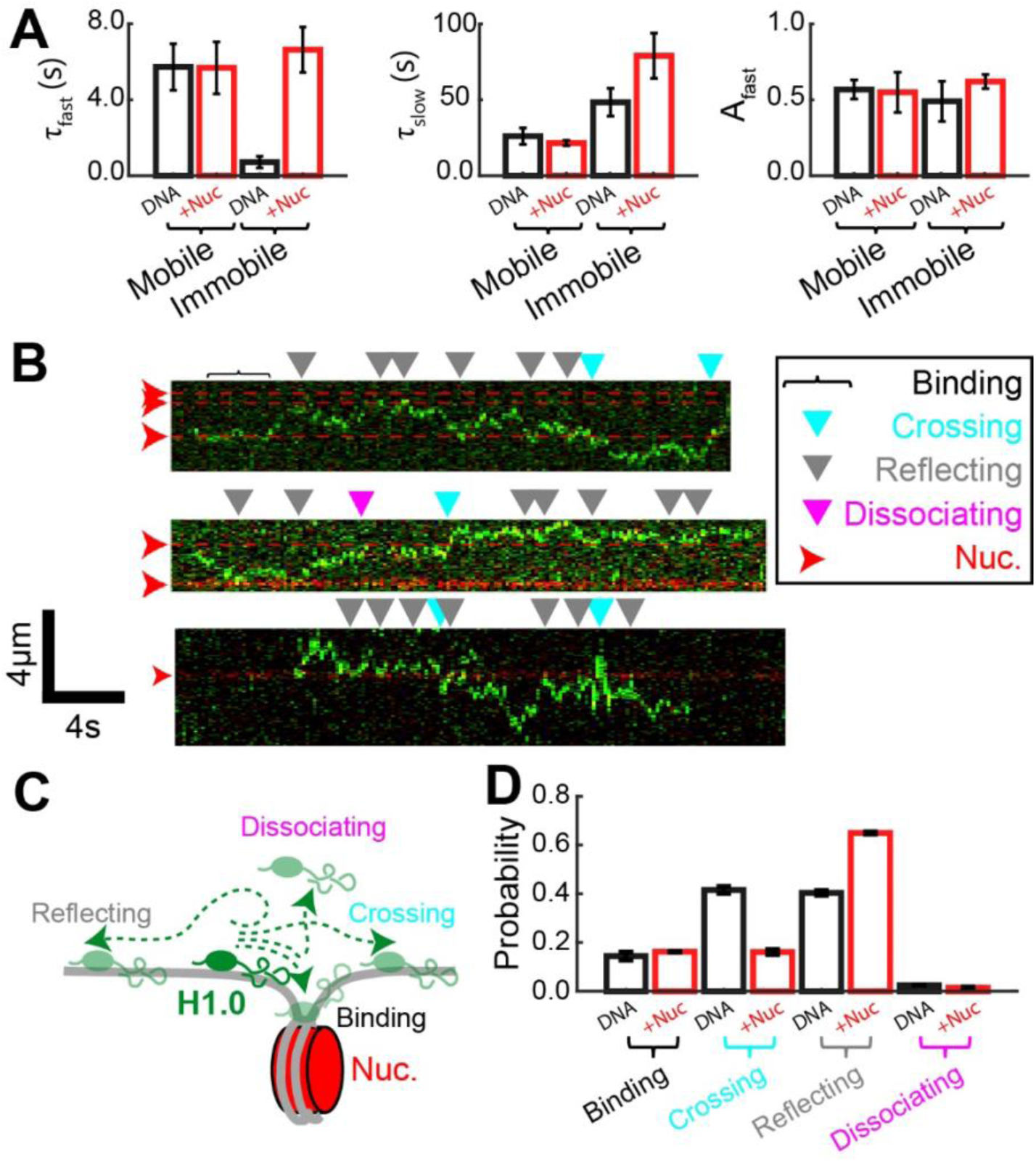
Summaries of H1.0 dynamics on nucleosome-containing tethers. (**A**) Fit parameters: τ_fast_, τ_slow_ and A_fast_ for CPDs of mobile and H1.0_immobile_ traces when bound to DNA-only (DNA, black) and nucleosomes-containing (+Nuc, red) tethers (Figure 2F-G). (**B**) Representative kymographs of H1.0_mobile_ (green) bound to nucleosome-containing tethers. The nucleosome locations are indicated by dotted red lines. (**C**) Illustration of the four possible outcomes following a colocalization event between a H1.0_mobile_ and nucleosome: binding, crossing, reflecting, and dissociating. (**D)** Summary of the probability of binding (0.4s < colocalization time), crossing, reflecting, and dissociating events for H1.0_mobile_-nucleosome colocalization events (red) and H1.0mobile colocalization events at randomly selected points along DNA-only tethers (black). All measurements were done with a constant force of 1 pN.

**Extended Figure 4.**
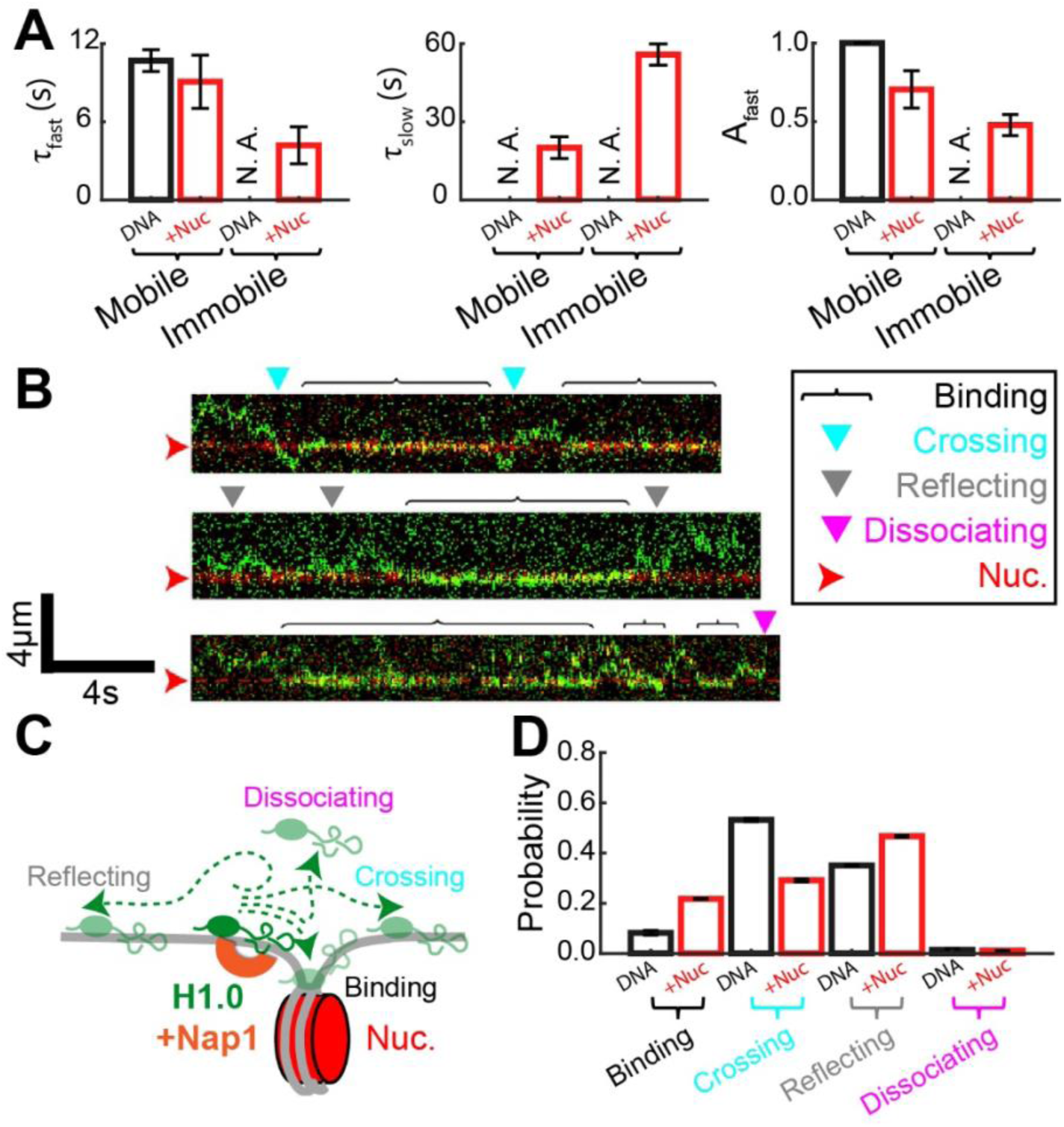
Summaries of H1.0 dynamics on nucleosome-containing tethers in the presence of Nap1. (**A**) In the presence of Nap1, fit parameters τ_fast_, τ_slow_ and A_fast_ for CPDs of H1.0_mobile_ and H1.0_immobile_ when bound to DNA-only (DNA, black) and nucleosomes-containing (+Nuc, red) tethers (Figure 4F-G). Nap1 eliminated the presence of H1.0_immobile_ on DNA-only tethers and is listed as not applicable (N. A.). The CPD of H1.0_mobile_ on DNA-only tethers fit best to a single exponential. Therefore, τ_slow_ is not applicable and A_slow_ = 1. (**B)** Representative kymographs of H1.0_mobile_ (green) bound to nucleosome containing tethers. The nucleosome locations are indicated by dotted red lines. (**C**) Illustration of the four possible outcomes following a colocalization event between a H1.0_mobile_ and nucleosome: binding, crossing, reflecting and dissociating. (**D)** Summary of the probability of binding (0.4s < colocalization time), crossing, reflecting, and dissociating events for H1.0_mobile_-nucleosome colocalization events (red) and H1.0_mobile_ colocalization events at randomly selected points along DNA-only tethers (black). All measurements were done with a constant force of 1 pN and 2nM Nap1.

